# Evaluating wildlife rehabilitation: Successful return of rehabilitated gulls in the wild

**DOI:** 10.1101/2025.09.22.677698

**Authors:** Tine Uytterschaut, Reinoud Allaert, Wouter Courtens, Claude Velter, Wendt Müller, Eric Stienen, An Martel, Luc Lens, Frederick Verbruggen

## Abstract

The expansion of human activities has increased the need for animal rehabilitation, especially for urban-breeding species such as herring gulls (*Larus argentatus*) and lesser black-backed gulls (*L. fuscus*), which now commonly nest on buildings and industrial sites. This study evaluates rehabilitation effectiveness by comparing survival and post-release reintegration, measured through habitat use and the degree of association with human-dominated environments, between rehabilitated and wild juvenile gulls along the Belgian coast. From 2012 to 2023, 3443 juveniles were ringed and monitored. Estimated survival rates varied significantly with age, but there was no difference between rehabilitated and wild birds. Differences in habitat use and association with local human population density were largely confined to juveniles: rehabilitated juveniles were resighted more often in rural green areas with lower human population densities, with patterns converging in older age classes. Our results suggest rehabilitation can support urban gull populations by yielding wild-like survival and space use.

## 1. Introduction

Human-induced environmental change, particularly the rapid expansion of urban areas, is transforming ecosystems worldwide, fundamentally reshaping species interactions and increasing encounters between humans and wildlife. Although some species seem to thrive in urban environments by exploiting new resources, others face habitat fragmentation, increased human-wildlife conflicts, and other hazards that can lead to injury, displacement, or population declines. Common urban threats include vehicle collisions, building strikes, and entanglement with human-made structures (Cope et al., 2022; Kwok et al., 2021; Miller et al., 2023; Montesdeoca et al., 2017; Souëf et al., 2015; Tribe and Brown, 2000). In response, wildlife rehabilitation centres provide temporary care for injured, sick, or orphaned animals with the goal of reintroducing them to the wild (*International Wildlife Rehabilitation Council* 2024). Each year, millions of animals undergo rehabilitation globally (Cope et al., 2022; Kwok et al., 2021), yet the long-term effectiveness of these efforts remains unclear. A first essential step is to assess post-release survival of rehabilitated animals to determine whether rehabilitation contributes meaningfully to species conservation, ensuring that resources are allocated to strategies with the greatest ecological impact (Pyke and Szabo, 2017). Equally, assessing post-release behaviour is necessary, as survival alone does not guarantee successful reintegration.

Although many rehabilitation centres report release rates of 40–55% (Hanson et al., 2021; Kwok et al., 2021; Montesdeoca et al., 2017; Mullineaux and Pawson, 2023), few studies have systematically tracked released animals to quantify post-release survival, and even fewer have examined how these individuals reintegrate behaviourally into wild populations (Pyke and Szabo, 2017). Studies tracking post-release survival report mixed outcomes. In species such as yellow-eyed penguins (*Megadyptes antipodes*) and cape vultures (*Gyps coprotheres*), rehabilitated individuals often exhibit lower survival than wild conspecifics (Alden et al., 2021; Monadjem et al., 2013), whereas peregrine falcons (*Falco peregrinus*), black bears (*Ursus americanus*), and Carnaby’s cockatoos (*Zanda latirostris*) often exhibit comparable survival to wild conspecifics (Blair et al., 2019; Groom et al., 2017; Sweeney et al., 1997). In some cases, both survival and behaviour of rehabilitated animals resemble those of wild conspecifics; for example, rehabilitated western gulls (*Larus occidentalis*) and little penguins (*Eudyptula minor*) have been shown to rejoin colonies and forage similarly to wild birds (Chilvers et al., 2015; Golightly et al., 2002). Other species may still exhibit shifts in behaviour. For example, harbour seals (*Phoca vitulina*) range more widely after release (Gaydos et al., 2012; Sangster et al., 2021), and brown pelicans (*Pelecanus occidentalis*) display altered dispersal distances and were less likely to attend breeding sites (Lamb et al., 2018). One explanation is that although rehabilitation centres provide veterinary care, nutrition, and protection from predators, captivity can impede the development of critical survival skills, such as efficient foraging, predator avoidance, and social interactions, that are potentially acquired through prolonged parental care in the wild (Ellis et al., 2000; Griffin, 2004; Kreger et al., 2005). Early separation and time in artificial conditions may therefore compromise competitive ability and reduce post-release survival across a range of rehabilitated species.

Against this background, gulls provide a useful case study. In Europe, herring gulls (*Larus argentatus*) and lesser black-backed gulls (*Larus fuscus*) have increasingly shifted from coastal to urban breeding sites, driven primarily by loss of natural breeding habitat through coastal development (e.g., port expansion), increased predation pressure on the ground, and the availability of reliable anthropogenic food sources such as landfill waste and urban refuse (Freed et al., 2021; Goumas et al., 2024; Kavelaars et al., 2020; Monaghan, 1979; Rock, 2005; Stewart et al., 2020). A key challenge of urban nesting arises when gull chicks, typically around four weeks of age, begin exploring their surroundings before they are fully capable of flight, a behaviour potentially linked to heat stress when rooftop surfaces become dangerously hot (pers. obs., wildlife rescue centre Ostend). Although falls rarely cause severe injury, chicks are frequently rescued and taken to rehabilitation centres, on average 149 per year during the study period, because returning them to nests on high, hard-to-access rooftops is often not feasible. In these centres, chicks are temporarily housed and cared for until they reach independence.

As a first step in evaluating rehabilitation success for these gulls, we compared survival of rehabilitated and wild birds using a subset of long-term ringing data from the Research Institute for Nature and Forest (INBO). Since 1986, INBO has maintained a long-term ringing programme of lesser black-backed and herring gulls, with extensive resightings reported across the northern Netherlands to southern Spain by researchers and citizen scientists. Using capture-mark-recapture models, we estimated survival probabilities across species and age classes as these gulls matured. In a second step, we contrasted post-release habitat use and the human population density at resighting locations to evaluate how rehabilitated juveniles behave compared to wild conspecifics.

We hypothesise that, due to the absence of parental guidance during critical developmental periods, rehabilitated juveniles will (i) exhibit lower survival rates than wild counterparts, (ii) differ in habitat selection, and (iii) show stronger associations with densely populated areas, reflecting their urban origin and expected tolerance of humans. Our study provides a comprehensive assessment of rehabilitation effectiveness, informing conservation strategies for large gulls.

## 2. Material and Methods

### (a) Data collection

We analysed colour-ring data of wild and rehabilitated herring gulls and lesser black-backed gulls ringed as juveniles between 2012 and 2019 to compare survival rates and habitat use. We excluded individuals ringed before 2012, as consistent colour-ringing at the rehabilitation centre only began that year. Birds ringed after 2019 were excluded, allowing a minimum of five years of resighting data for each individual to reach adulthood. All ringing was conducted under INBO and Ghent University licences. Each bird was fitted with a standard metal ring and a uniquely coded plastic colour ring (blue with white alphanumeric characters). Resightings were reported opportunistically by researchers and the public, with variable observer effort and no standardised survey protocol. Our dataset comprises two species, each divided into two groups: herring gulls (n = 1,537; rehabilitated = 711, wild = 826) and lesser black-backed gulls (n = 1,907; rehabilitated = 389, wild = 1,517). Rehabilitated birds were treated at the Wildlife Rehabilitation Centre of Ostend and ringed prior to release, while wild birds were ringed as 3–5-week-old chicks in the Ostend colony (7 km from the rehabilitation centre) and the Zeebrugge colony (28 km). At the centre, these predominantly uninjured juveniles were housed in large outdoor flight cages and raised in mixed groups of herring and lesser black-backed gulls until they were capable of sustained flight, after which they were released. To avoid confounding factors, we omitted 31 individuals involved in experimental studies and six fitted with tracking devices. We excluded 25 wild chicks that never fledged and 11 rehabilitated chicks that were not released after ringing. The final database included 3,443 ringed individuals. Annual and species-specific ringing totals are provided in Supplementary Table S1.

#### (i) Survival dataset preprocessing

Live resightings of ringed birds, collected between July 2012 and November 2023 by researchers and citizen scientists, were compiled into annual encounter histories for each bird, with “1” indicating an alive observation and “0” otherwise, following the Cormack–Jolly–Seber (CJS) framework (Lebreton et al., 1992). The CJS framework was chosen for its efficiency in estimating apparent survival and detection probabilities from binary encounter histories under moderate data requirements. We restricted resighting records to July–November, the period with highest reporting rates and aligned with wild chicks’ fledging in July, following the method of Kentie et al., 2022. In addition, we limited the geographic scope of the study to areas with the highest resighting effort (50°N–53,5°N, 1°W–6°E; Figure 1).

**Figure 1.**
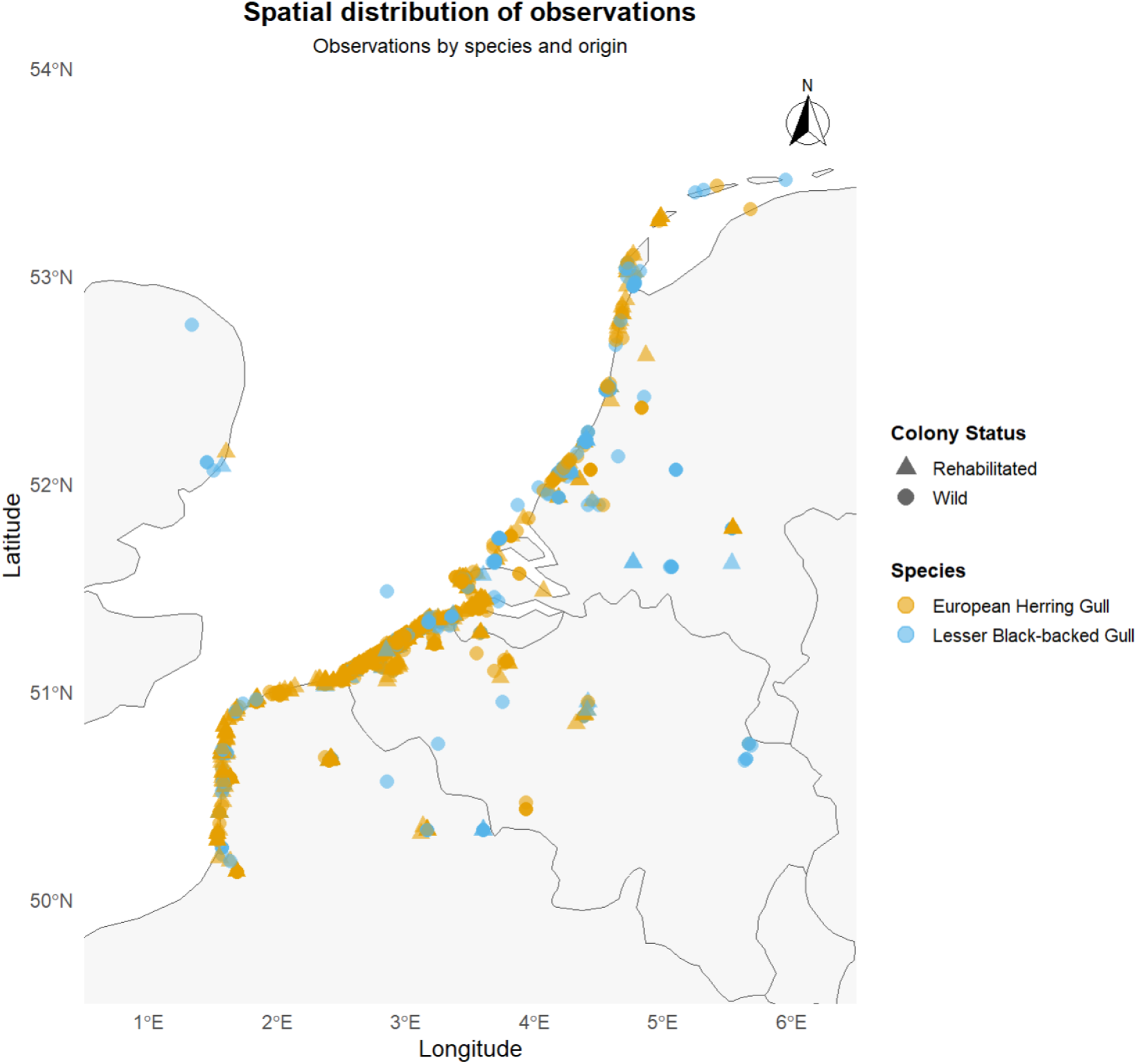
Spatial distribution of the selected resightings.

The final dataset comprised 22,733 observations from 3,443 individuals (2,343 wild; 1100 rehabilitated). To account for the effect of transience, and to capture major life-history stages, while ensuring sufficient sample sizes per class, birds were assigned to three age classes: juvenile (year 1), immature (years 2–4) and adult (year 5+). We opted for this approach as continuous-age models or finely stratified multistate approaches would require complex parametric or numerous age-specific parameters, potentially leading to overfitting, imprecise estimates, or convergence issues given sparse age-specific resightings.

#### (ii) Habitat use and population density preprocessing

For the habitat use and population density analyses, we compiled all available resightings of the selected juveniles from 2012 to 2023, irrespective of season or location, to characterize post-fledging space use. To ensure that the data reflected meaningful post-fledging habitat use, we applied two filters. First, we excluded ringing records within the colony boundaries, as these do not represent free-ranging behaviour (see Supplementary Figure 5). Second, we excluded wild-juvenile observations that occurred before the first release date of rehabilitated juveniles in the same year (245 observations), thereby aligning the observation window across groups. The final dataset included 31,623 observations from 2,488 individual gulls, comprising 1,388 wild and 1,100 rehabilitated individuals.

Next, each observation was annotated using the 2018 reference year of the European Corine Land Cover inventory (100m resolution; Service, 2020). We consolidated the original 44 land cover classes into seven main habitat types: (1) intertidal areas, (2) rural green areas, (3) built-up areas, (4) port areas, (5) city green areas, (6) industrial areas, and (7) waste processing areas, broadly following (Spelt et al., 2019) (see Supplementary Table S7 for details). Because coordinate precision varied among years, with older records often rounded to only two decimal places, some locations were imprecise. We therefore recoded habitat values based on the accompanying place names (see Supplementary Table S8). To identify nest sites, we overlaid a custom layer of polygons around known colonies in Ostend and the harbour of Zeebrugge; observations within these were labelled “natural nestsite.” Observations on the rehabilitation centre roof were labelled “artificial nestsite” (see Supplementary Figure 5). Both nestsite categories were excluded from all habitat-use analyses.

To explore the relationship between gull resightings and human presence, we extracted population density values from the 2020 Global Human Settlement Layer (Copernicus, 1000m resolution (Pesaresi et al., 2024)), using bilinear interpolation. Missing population density values were replaced with zero, and the response variable (plus one) was log-transformed to reduce skewness.

### (b) Statistical analyses

#### (i) Survival analysis

Survival probability (*Φ*) and detection probability (*p*) were estimated using CJS models (Cormack, 1964; Jolly, 1965; Seber, 1965) in RStudio (version 2025.05.01+513, with R version 4.5.0) with the RMark package (Laake, 2013, version 3.0.0). Estimates were generated for three age classes: Juvenile (1 year), Immature (2-4 years), and Adult (*≥*5 years), following Kentie et al. (2022). Survival was modelled as a function of age class, species, and prefledging status (wild vs. rehabilitated), while detection probability also included time as a covariate. Model selection was conducted in two steps to manage the large number of possible combinations for *Φ* and *p*. In the first step, we fixed the most complex survival structure and compared multiple detection models (Supplementary Table S3). Goodness-of-fit (GOF) tests were performed in program MARK (White and Burnham, 1999) using a parametric bootstrap (n = 100), yielding a variance inflation factor (ĉ) of 1.257, indicating no substantial overdispersion (Burnham and DR Anderson, 2002)^1^. Model selection was based on the QAIC_*c*_, adjusted for small sample size and ĉ. The top-ranked detection model included an interaction between species and age class, along with main effects of prefledging status and time. In the second step, we fixed this best-fitting *p*-structure and evaluated all combinations of *Φ* (Supplementary Table S4). GOF testing for the most complex survival model yielded a ĉ of 1.230, again suggesting no overdispersion. The final CJS model retained interactions between prefledging status and species, and between species and age class. Separate species-specific models (see Supplementary Table S2) yielded the same top-ranked structures, supporting our choice to present the combined two-species analysis for clarity and statistical robustness.

#### (ii) Habitat use analysis

While survival was analysed in a combined two-species framework, habitat use was modelled separately for each species to avoid an excessive number of interaction terms and to improve model interpretability. We used generalised linear models with a negative binomial distribution in the glmmTMB package (Brooks et al., 2017). For each species, we modelled resighting counts, grouped by habitat category, as a function of a three-way interaction among prefledging status (wild vs. rehabilitated), age class (juvenile, immature, adult), and habitat classification. To account for variation in overall observation effort, the total number of resightings per individual was included as an offset (log-transformed). We evaluated model fit and residual structure using the `performance` (Lüdecke et al., 2019) and `DHARMa` (Hartig, 2016) packages. Predicted resighting counts on the response scale were extracted for every combination of prefledging status, age class and habitat using emmeans (Lenth, 2025). Differences between rehabilitated and wild birds within each habitat–age combination were assessed using pairwise contrasts from the emmeans package, with a Holm adjustment applied to control the family-wise error rate, as only planned comparisons between rehabilitated and wild birds were conducted.

#### (iii) Population density analysis

To test whether human population density at sighting locations differed by prefledging status, we fitted zero-inflated generalised linear mixed models (ZIGLMMs) via glmmTMB (Brooks et al., 2017). A zero-inflated approach was chosen because many resightings occurred at locations with a recorded human population density of zero, resulting in an excess of zeros in the data. These models comprise two components: a conditional component, which models the (transformed) population density where observations occur, and a zero-inflation component, which models the probability of structural zeros, thus accounting for excess zeros and potential overdispersion (Zeileis et al., 2008). The initial conditional model included interactions between prefledging status and both age class and species, as well as a random intercept for individual identity. The interaction between prefledging status and species was removed due to lack of significance (*p* = 0.999), yielding a final conditional model with a prefledging status × age class interaction and a main effect of species. The zero-inflation submodel initially included all two-way interactions among prefledging status, age class, and species. After comparing AIC values, we removed the the prefledging status x age class interaction and the prefledging status main effect (both non-significant and collinear with species), retaining only species and age class as zero-inflation predictors. Model fit and residual diagnostics were assessed using the DHARMa package (Hartig, 2016), confirming adequate fit and no remaining overdispersion. Marginal means and pairwise contrasts between rehabilitated and wild groups within each age class and species were extracted using the emmeans package (Lenth, 2025), with a Tukey adjustment applied to control the family-wise error rate across all pairwise comparisons among factor levels.

## 3. Results

### (a) Survival

The best-fitting model indicated that apparent survival varied depending on prefledging status, species, and age class (Figure 2, Supplementary Table S5). Overall, rehabilitated and wild individuals exhibited similar survival probabilities. However, a model including a prefledging status × species interaction fitted significantly better than one without the interaction term (*χ*^2^ = 6.02, df = 1, *p* = 0.0142), suggesting that survival probabilities differed by species. Wild lesser black-backed gulls showed slightly higher survival probabilities than rehabilitated individuals (Figure 2, Supplementary Table S5), although this contrast across age classes was not statistically significant in post-hoc pairwise comparison (*p* = 0.241). No such numerical difference was observed for herring gulls.

**Figure 2.**
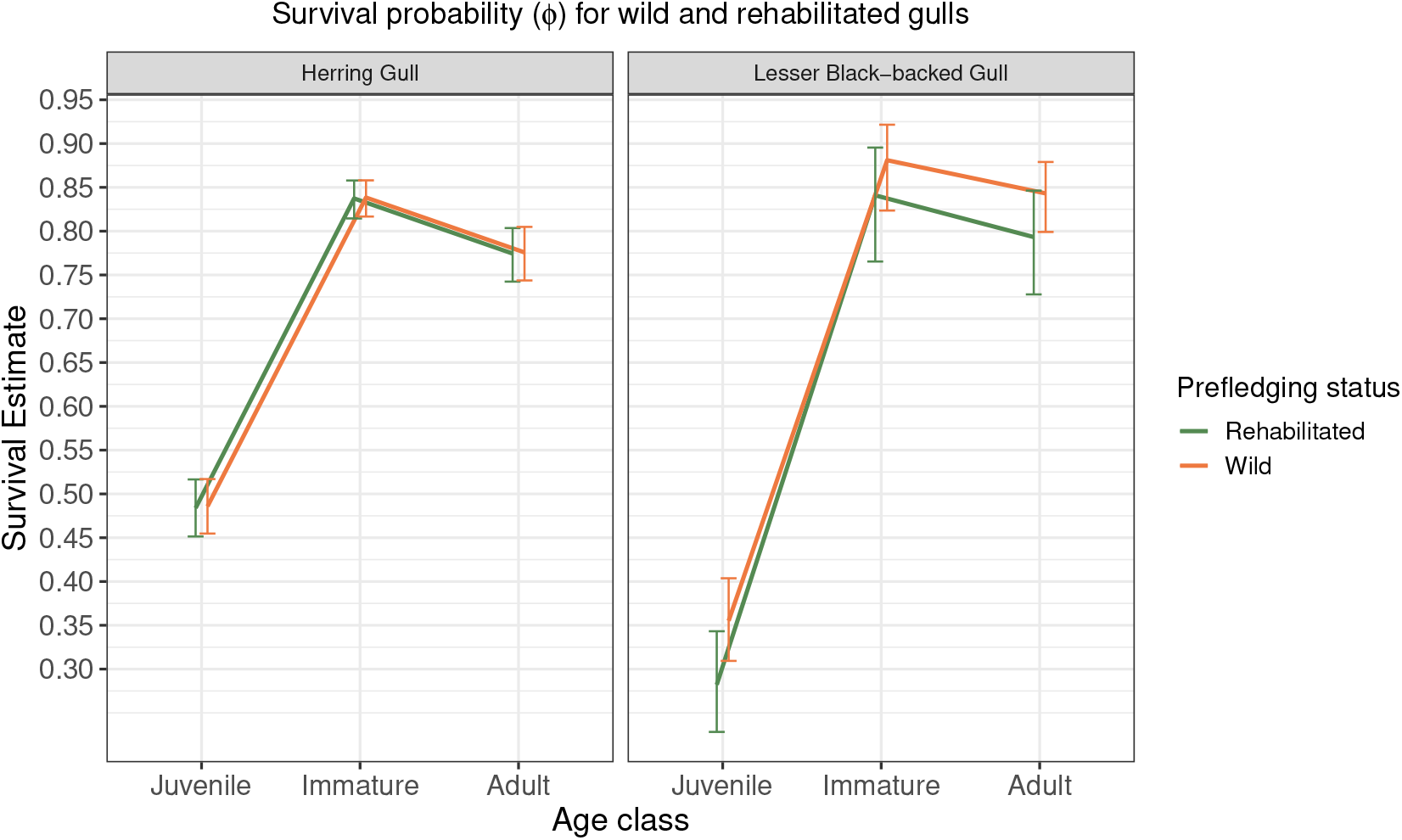
Estimated apparent survival (*ϕ*) for wild (red) and rehabilitated (green) herring gulls (left) and lesser black-backed gulls (right) for juveniles (age 1), immatures (age 2-4) and adults (age 5+).

Survival trajectories increased from the juvenile to the immature stage and declined slightly in adulthood, with largely overlapping estimates between wild and rehabilitated birds (Table S6). For herring gulls, estimated juvenile survival was virtually identical between wild (48.6%, 95% CI: 0.46–0.52) and rehabilitated birds (48.4%, 0.45–0.52). In the immature stage, survival rose to 83.8% (0.82–0.86) in wild and 83.7% (0.82–0.86) in rehabilitated individuals. Adult survival was slightly lower, at 77.6% (0.74–0.81) for wild and 77.4% (0.74–0.80) for rehabilitated birds. In lesser black-backed gulls, juvenile survival was lower overall and showed more numerical divergence between groups, with 35.5% (0.31–0.40) in wild and 28.2% (0.23–0.34) in rehabilitated birds. However, these differences diminished in later stages: immature survival reached 88.1% (0.82–0.92) for wild and 84.1% (0.77–0.90) for rehabilitated individuals, while adult survival was 84.3% (0.80–0.88) and 79.3% (0.73–0.85), respectively.

Age-class effects differed between species, as a model including a species × age class interaction in survival fitted significantly better than the additive model (likelihood ratio test: (*χ*^2^ = 24.39, df = 2, *p <* 0.001)), indicating a stronger juvenile-to-immature and juvenile-to-adult survival gain in lesser black-backed gulls compared to herring gulls. However, post-hoc pairwise contrasts of absolute survival in the immature (*p* = 0.948) and adult (*p* = 0.718) stages revealed no significant inter-specific differences.

### (b) Habitat use

Resighting distributions across habitat types differed between wild and rehabilitated gulls (Fig. 3). Estimated marginal means (on the response scale) are reported in the text, while relative risk ratios between groups are presented in Tab. S9. For herring gulls, differences between wild and rehabilitated birds occurred in four habitat–age combinations. First, rehabilitated juveniles were resighted more often in intertidal areas (8.703 *±* 0.386 SE vs. 6.877 *±* 0.410 SE; *p <* 0.01). Second, among immatures, rehabilitated birds were more frequently resighted in industrial areas (1.700 *±* 0.386 SE vs. 0.668 *±* 0.152 SE; *p <* 0.01). Third, in adults, rehabilitated birds were more frequently resighted in industrial areas (1.735 *±* 0.375 SE vs. 0.834 *±* 0.153 SE; *p <* 0.001). Fourth, rehabilitated adults were more frequently resighted in waste processing areas (3.610 *±* 0.422 SE vs. 2.414 *±* 0.284 SE; *p* = 0.015). No other habitats differed by origin in any age class (all *p >* 0.05).

**Figure 3.**
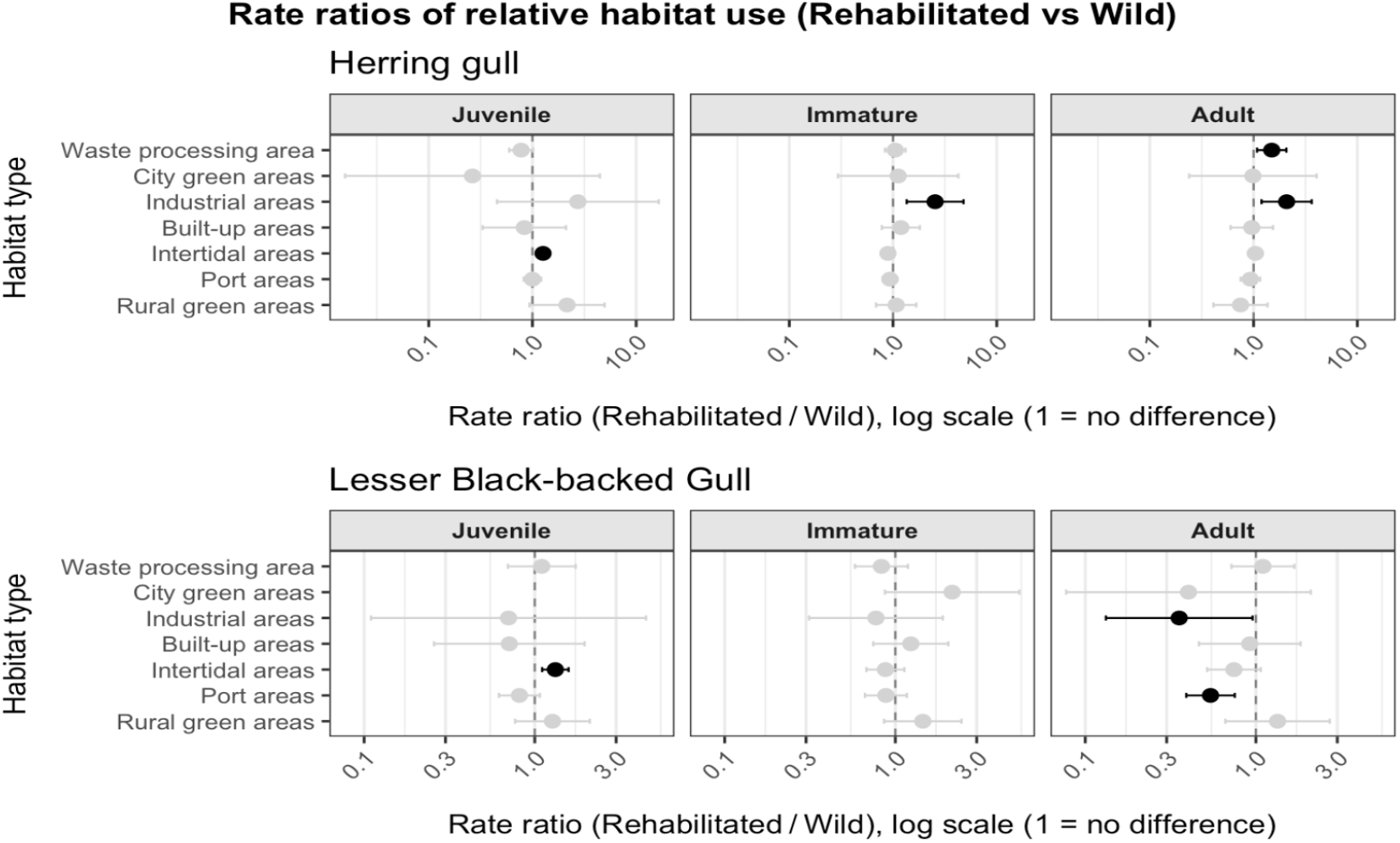
Forest plots of rate ratios (Rehabilitated vs. Wild) for habitat use in herring gulls and lesser black-backed gulls by age class. Points show the estimated incidence-rate ratio of resightings in each habitat per unit effort, with horizontal whiskers indicating 95% confidence intervals. Comparisons for which the interval does not include 1 (i.e. significant differences) are shown in black. The vertical dashed line at 1 indicates no difference between rehabilitated and wild birds.

For lesser black-backed gulls, origin-related differences were confined to intertidal, industrial, and port areas and varied with age (Fig. 3, Tab. S10). Rehabilitated juveniles were resighted more often in intertidal areas (4.050 *±* 0.237 SE vs. 3.067 *±* 0.216 SE; *p <* 0.01). Among adults, rehabilitated birds were less frequently resighted in industrial areas (0.677 *±* 0.326 SE vs. 1.906 *±* 0.295 SE; *p* = 0.04) and port areas (2.128 *±* 0.332 SE vs. 3.924 *±* 0.233 SE; *p <* 0.01). No other habitats differed by origin in any age class (all *p >* 0.05). Juvenile city green areas were rare and non-estimable for rehabilitated birds.

### (c) Population density

Across both species, rehabilitated juveniles were observed in locations with substantially lower human population density than wild juveniles (herring gull: 347.2 [323.2–372.9] vs. 899.9 [832.5–972.8] people km^2^; lesser black-backed gull: 260.0 [238.3–283.8] vs. 674.4 [627.1– 725.3] people km^2^; both *p <* 0.001). From the immature stage onward, densities converged as rehabilitated birds shifted into more populated areas while wild birds dispersed into lower-density zones. By the immature and adult stages, both groups occupied sites of comparable human population density (Table S12, Figure 4).

**Figure 4.**
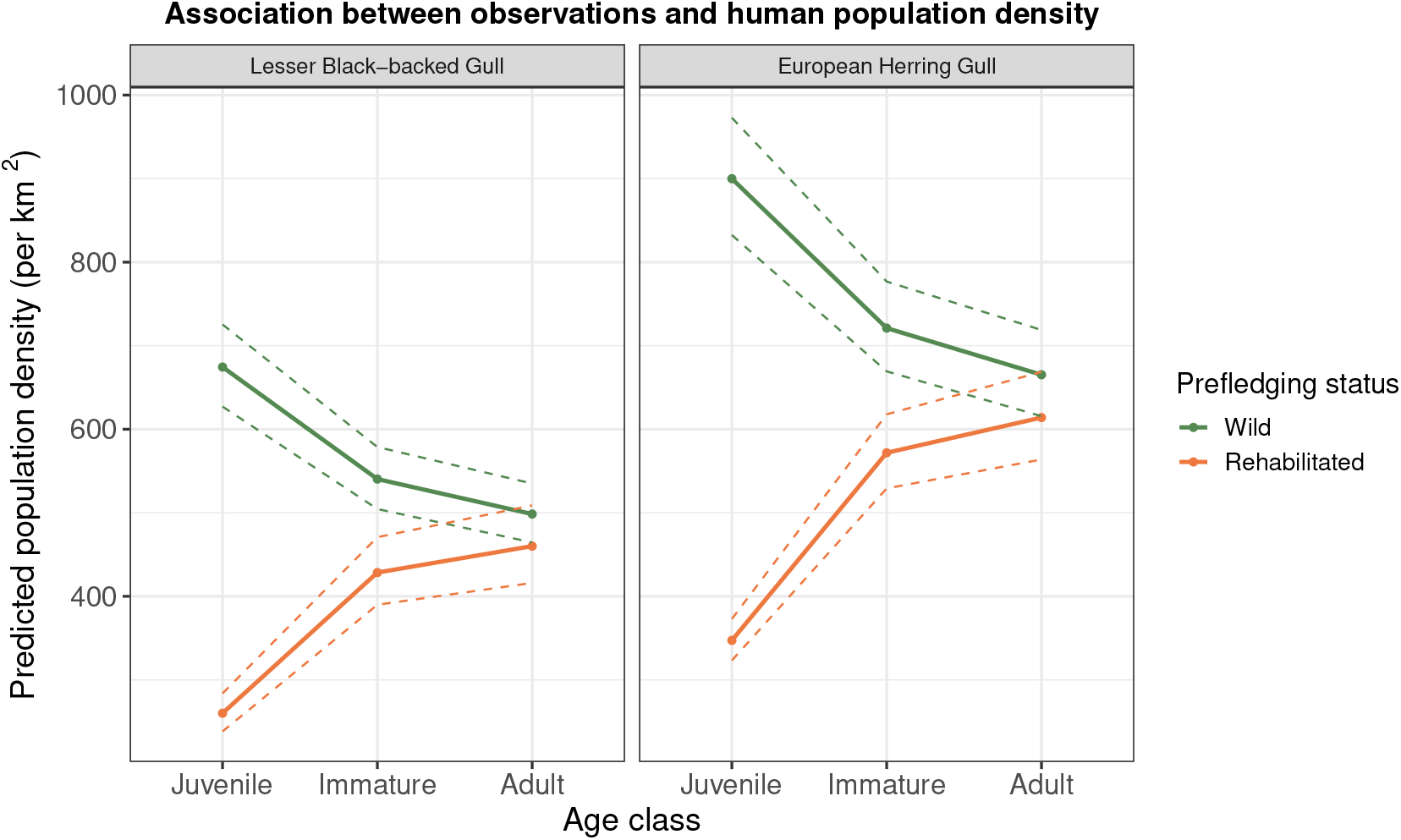
The association between observations and human population density by age class and species. The lines represent the predicted human population density per *km*^2^ where the gulls are observed, while the dashed lines indicates the 95% confidence intervals.

**Figure 5.**
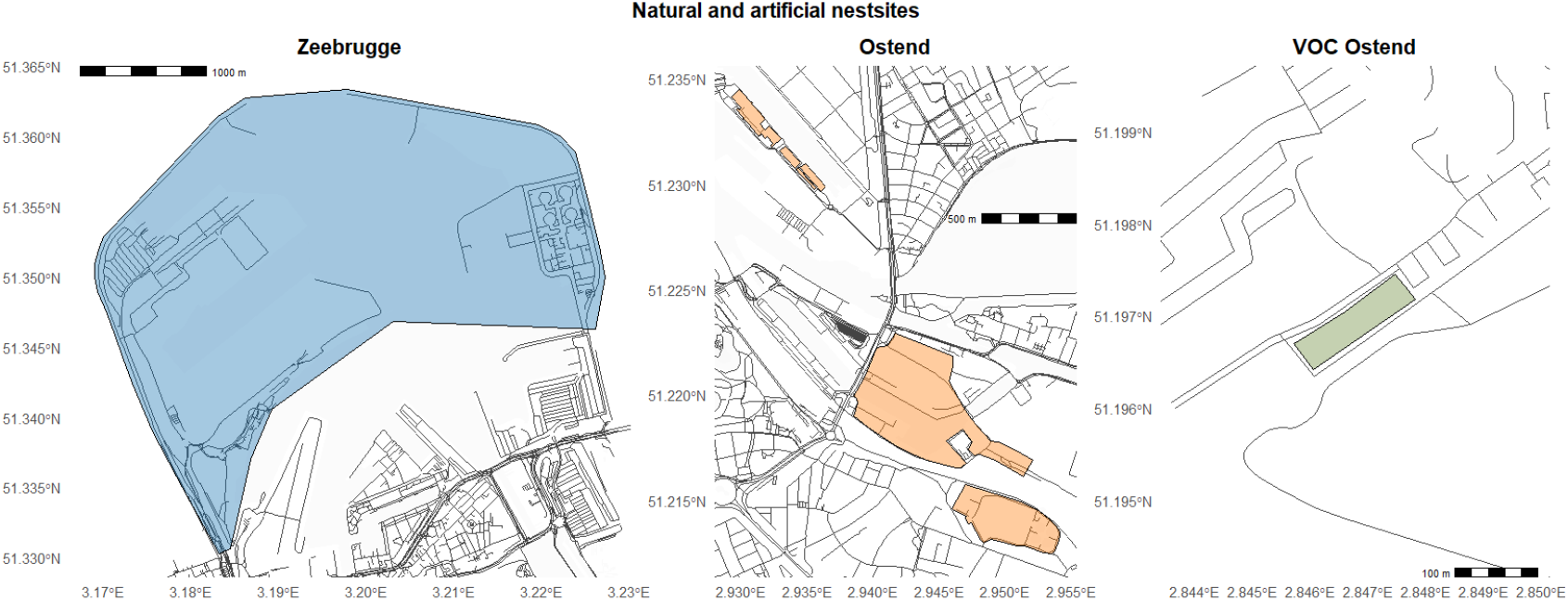
Natural nesting areas in Zeebrugge (left) and Ostend (right) and the rehabilitation centre as artificial nestsite.

**Figure 6.**
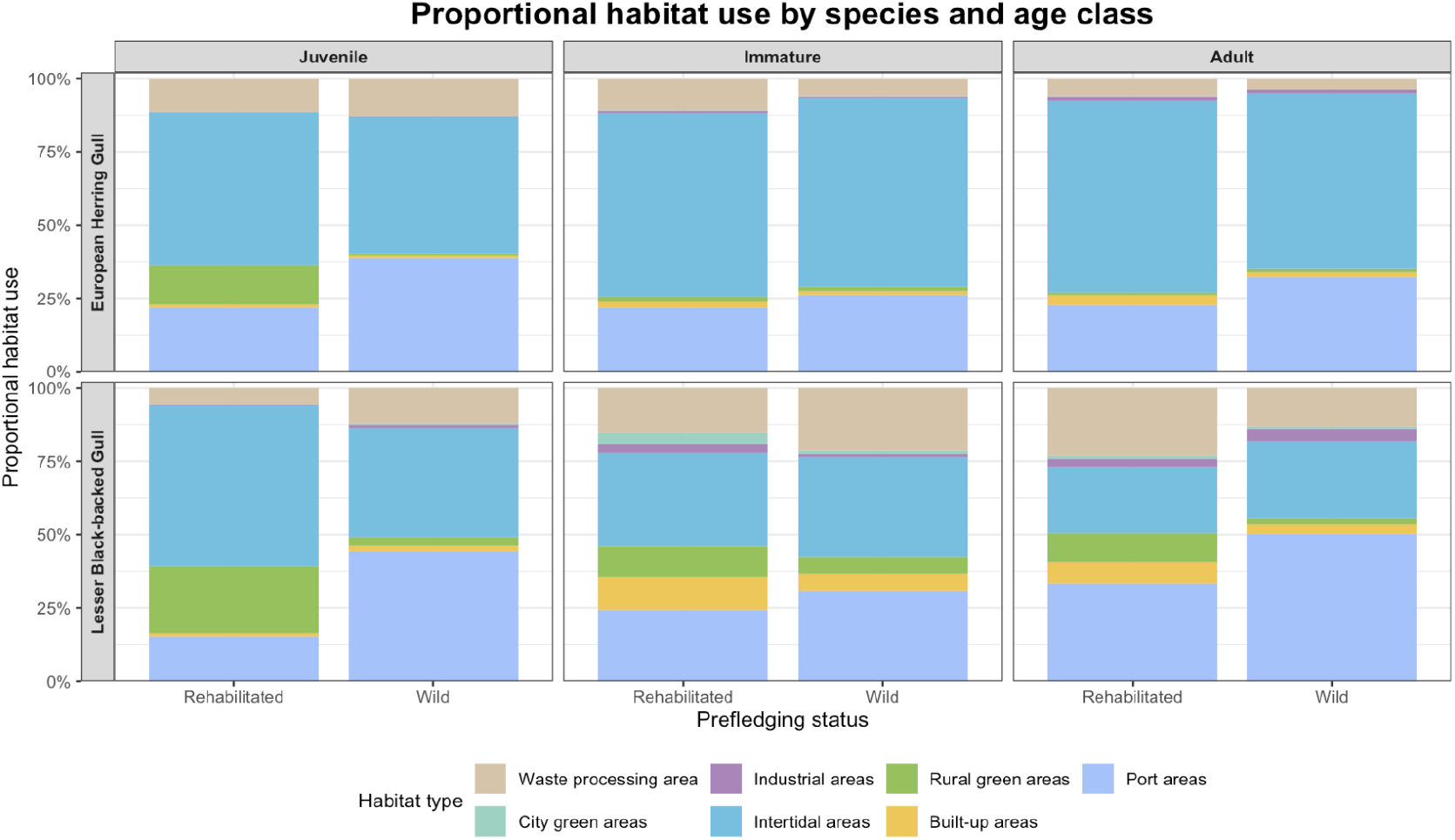
Proportional habitat use for herring gulls (top) and lesser black-backed gulls (bottom) across prefledging status and age classes. Proportions were calculated within each age class and prefledging status by normalizing predicted counts across habitat types

## 4. Discussion

By comparing the apparent survival probabilities of rehabilitated juvenile gulls with those of wild conspecifics, we found no evidence that release from rehabilitation incurs a lasting survival penalty. Survival in the first year, immature stage, and adulthood did not differ significantly between groups, contradicting our initial expectation that rehabilitated juveniles would fare worse. This suggests that, despite potential early-life deficits, such as reduced foraging proficiency or predator-avoidance skills (Alden et al., 2021; Monadjem et al., 2013), rehabilitated juveniles attained survival comparable to wild-reared counterparts over the period monitored. Behavioural patterns inferred from resighting data showed age-dependent nuances: as juveniles, rehabilitated birds of both species were more frequently resighted in intertidal areas; in the immature stage, rehabilitated herring gulls were more often observed in industrial areas, whereas no consistent differences were detected in lesser black-backed gulls; and in adulthood, rehabilitated herring gulls were more frequently resighted in industrial areas and at waste-processing sites, while rehabilitated lesser black-backed gulls were less often observed in industrial and port habitats. No other consistent habitat differences emerged between origins. In addition, rehabilitated juveniles of both species were initially recorded in areas of lower human population density but converged with wild birds by adulthood.

Survival trajectories for both wild and rehabilitated gulls followed the typical life-history pattern of long-lived seabirds: low first-year survival, a marked increase during the immature stage, and a modest decline in adults. Previous studies reported that first-year survival in large gulls generally ranges from 30–70% in juveniles, increasing to 80–90% in adults, with considerable variation among years and colonies (Camphuysen and Gronert, 2012; Kentie et al., 2022; Rock and Vaughan, 2013; Schekkerman et al., 2021). Our estimates (≈35–50% for juveniles; ≈80–85% for adults) fall within these published ranges. Although our comparisons revealed no significant differences between origins, it is important to note that even small differences in early survival can translate into large cumulative losses over a long lifespan (Lebreton et al., 1992). Thus, non-significant trends should be interpreted cautiously in the context of population-level consequences.

Once fledged, rehabilitated and wild-reared juveniles exhibited indistinguishable age-structured survival patterns, with statistically equivalent annual survival in the immature and adult stages. The slight decrease from the immature to adult stage is probably associated with the costs of first-time breeding, rather than senescence or ring loss, as has been suggested in other studies. This is because our study period is shorter than both species’ lifespans and the longevity of plastic rings (Allen et al., 2019; Reed et al., 2008).

An inter-species comparison revealed that juvenile lesser black-backed gulls had lower apparent survival (34.9%) than juvenile herring gulls (49.6%), while survival converged in later stages (85.8% and 84.7% in immatures, respectively). This difference at the juvenile stage is consistent with their contrasting post-fledging dispersal ecologies: juvenile lesser black-backed gulls commonly undertake long-distance migrations to wintering grounds in southwestern Europe and Africa, whereas herring gulls in our study system remain largely resident or disperse only locally (Bustnes et al., 2013; Klaassen et al., 2012). However, the CJS models used here cannot distinguish true mortality from permanent emigration. Thus, individuals moving beyond our geographic scope may be misclassified as dead until they return. This well-known confounding of emigration with mortality likely depresses first-year survival estimates and exaggerates the apparent gain at the immature stage (Seber, 1965). Although genuine migratory mortality during the first migration also contributes (Oppel et al., 2015; Rotics et al., 2016), the observed pattern for lesser black-backed gulls therefore probably reflects a combination of true losses and undetected dispersal. Consequently, first-year survival is likely underestimated in this species, while immature survival is correspondingly overestimated due to returning emigrants. By adulthood, survival probabilities converge (79.5% for lesser black-backed gulls; 78.6% for herring gulls), indicating these early-life differences are transient. Crucially, this interaction does not alter our main conclusion: rehabilitation has minimal long-term impact on survival, as no comparisons between rehabilitated and wild cohorts were significant at any age class.

Our data indicate that rehabilitation does not appear to have long-term effects on survival, as rehabilitated juveniles showed outcomes comparable to wild-reared peers. However, most of the birds in our study were uninjured chicks rescued from city streets after early fledging, with limited flight capability leaving them vulnerable in high-traffic areas. Their comparable post-release survival likely reflects the benign nature of these admissions and may not extend to individuals released after severe injury or illness. Previous studies show that rehabilitation outcomes across taxa range from survival similar to wild conspecifics (DW Anderson et al., 1996; Duerr et al., 2023; Hagen et al., 2024; Punch, 2001) to markedly lower post-release survival (Alden et al., 2021; Barker et al., 2004; De La Cruz et al., 2013; Monadjem et al., 2013; Sharp, 1996), depending on cause and severity. This variation suggests that rehabilitation success is not only species-specific but also strongly influenced by admission length, type, and severity, which were not assessed here.

Beyond survival outcomes, our results also provide insight into post-release behaviour through patterns of habitat use. Rehabilitated juveniles of both species were disproportionately resighted in intertidal areas, characterized by lower competition and are easier for inexperienced birds to exploit, as more experienced adults tend to dominate higher-quality sites (Greig et al., 1983; Inzani et al., 2024; Monaghan, 1980; Spelt et al., 2019). Independent of habitat categories, rehabilitated juveniles also occurred in areas of lower human population density than wild juveniles, indicating a broader tendency to use less densely populated environments. The absence of extended post-fledging parental care, which in other bird species is known to increase juvenile survival (Grüebler and Naef-Daenzer, 2010; López-Idiáquez et al., 2018), likely exacerbates this constraint; wild juveniles are often still provisioned by their parents after fledging, allowing them to remain in higher-competition areas without having to rely (solely) on their own foraging skills (Burger, 1981). Over time, habitat use between the two groups converged, as wild birds dispersed into less populated areas and rehabilitated birds increasingly appeared in urbanised landscapes. Among adults, rehabilitated lesser black-backed gulls were under-represented at industrial and port habitats, reflected in the virtual absence of breeding records for rehabilitated individuals in these colonies. As these sites constitute the main monitored breeding areas of the species in our study area, this suggests that rehabilitated birds may breed elsewhere. In herring gulls, no such difference was detected; however, because this species frequently nests on rooftops, which are rarely monitored, comparable patterns may have gone unnoticed. Some of the observed differences, particularly the greater use of industrial and waste-processing sites by rehabilitated herring gulls at the adult stage, are difficult to interpret, as the activities underlying these resightings remain unclear. Addressing these uncertainties, and enabling direct comparisons of survival and habitat use, requires targeted GPS-tracking studies (Allaert et al. in prep).

Our inferences are based on opportunistic ring-resighting data, which are inherently biased towards locations with high observer activity (Thorup et al., 2014). Such bias can lead to overrepresentation of habitats near human population centres and underrepresentation of more remote areas, complicating interpretation of age- or species-specific patterns. Nevertheless, the observed differences are unlikely to be artefacts of detection bias. We accounted for spatial variation in observation effort by including a log-offset for individual observer effort and assume that any residual bias was comparable between rehabilitated and wild birds. Consequently, our findings should be interpreted as reflecting relative differences in habitat use attributable to prefledging status, rather than absolute habitat preferences. However, differences in habitat use persisted into adulthood, suggesting that rehabilitation may have longer-term behavioural consequences. Rehabilitated birds may breed at sites outside our monitoring, such as rooftops with lower hatching success (Van Malderen et al., unpublished). Together, our findings indicate that gull rehabilitation can yield survival rates comparable to those of wild conspecifics, while behavioural differences highlight the need for long-term monitoring. More broadly, placing equal emphasis on both survival and behavioural integration is essential for assessing the conservation and welfare value of rehabilitation across species.

## Author Contributions (CREDIT)

Conceptualization: T.U., R.A., L.L., F.V.; Methodology: T.U., R.A., W.C.; Data curation: T.U., R.A.; Formal analysis: T.U., R.A.; Visualization: T.U., R.A.; Validation: T.U., R.A., W.C.; Writing – original draft: T.U., R.A.; Writing – review and editing: T.U., R.A., W.C., C.V., W.M., E.S., A.M., L.L., F.V.; Supervision: L.L., F.V.; Funding acquisition: A.M., L.L., F.V.; Project administration: F.V.

## Acknowledgments

We are grateful to Marc Van De Walle, Hilbran Verstraeten, Nicolas Vanermen, and all colleagues at INBO for their extensive efforts in ringing. We also thank Hans Matheve (Ghent University), Isabelle Allemeersch, and Rijn Van Maele (Wildlife Rescue Centre) for their support. Over the years, more than 600 persons and organisations contributed to ring resightings and recoveries of these juveniles. We are especially grateful to Nathalie Colpaert, Roland François, Francis Kerckhof, Maarten van Kleinwee, Harry Vercruijsse, Jan Talloen, and Patrick Vandenbroucke for their dedicated contributions.

## Data, scripts, code, and supplementary information availability

All data required to replicate this study’s findings will be made openly available on Zenodo: https://doi.org/10.5281/zenodo.17078993. All scripts, code and raw data are accessible through the article’s OSF repository: https://osf.io/whczg/. Supplementary information supporting the results is also provided in this repository.

## Supplementary material

**Table S1.**
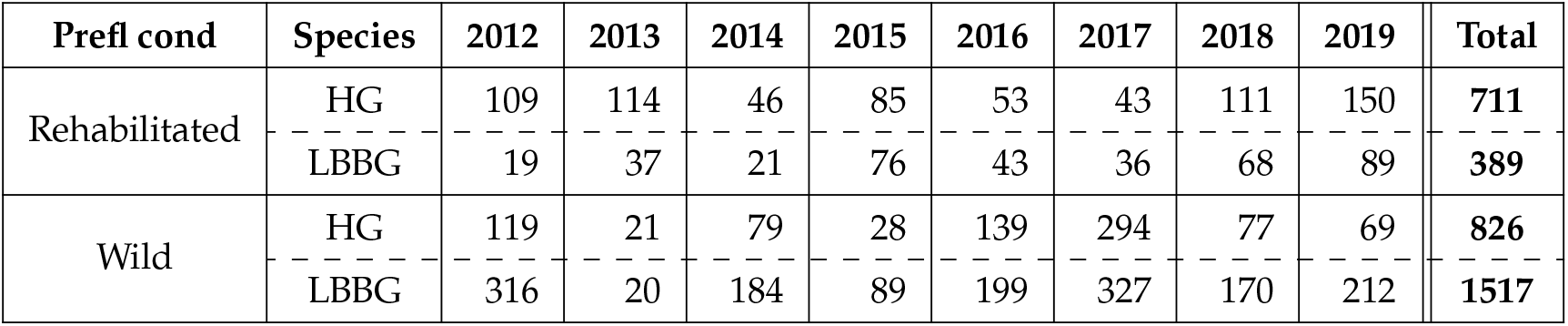
The number of rehabilitated and wild juvenile herring and lesser black-backed gulls ringed each year (2012-2019)

**Table S2.**
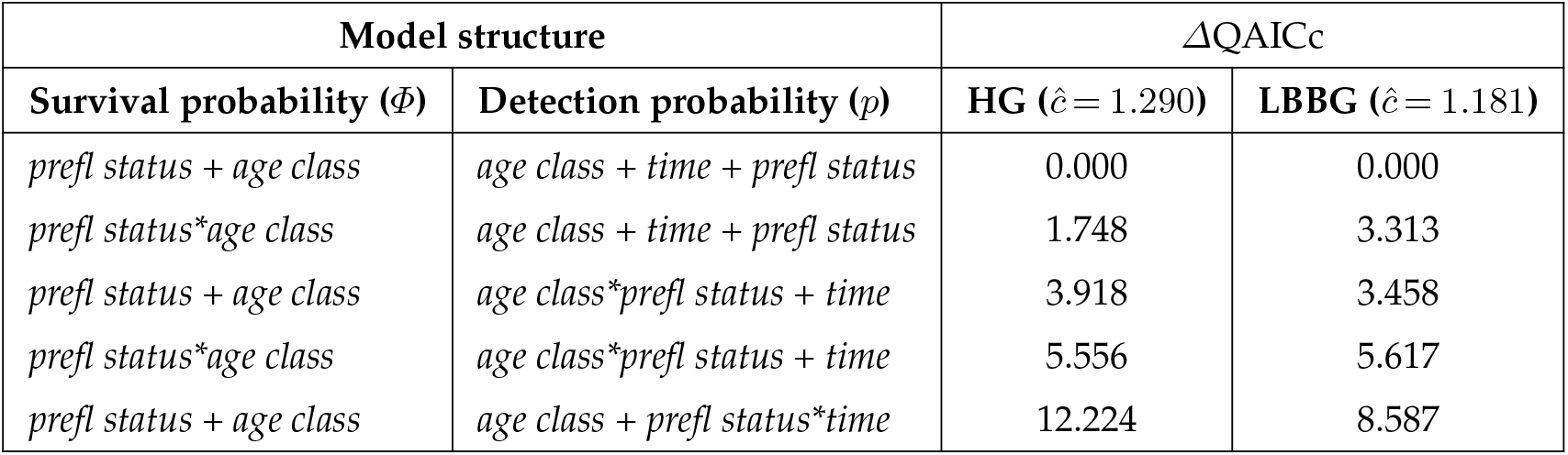
Top five ranked species-specific CJS models for herring gull (HG) and lesser black-backed gull (LBBG). Despite being modeled separately, the same structure ranked highest for both species, justifying the combined two-species analysis presented in the main text. *ĉ* values: HG = 1.290, LBBG = 1.181.

**Table S3.**
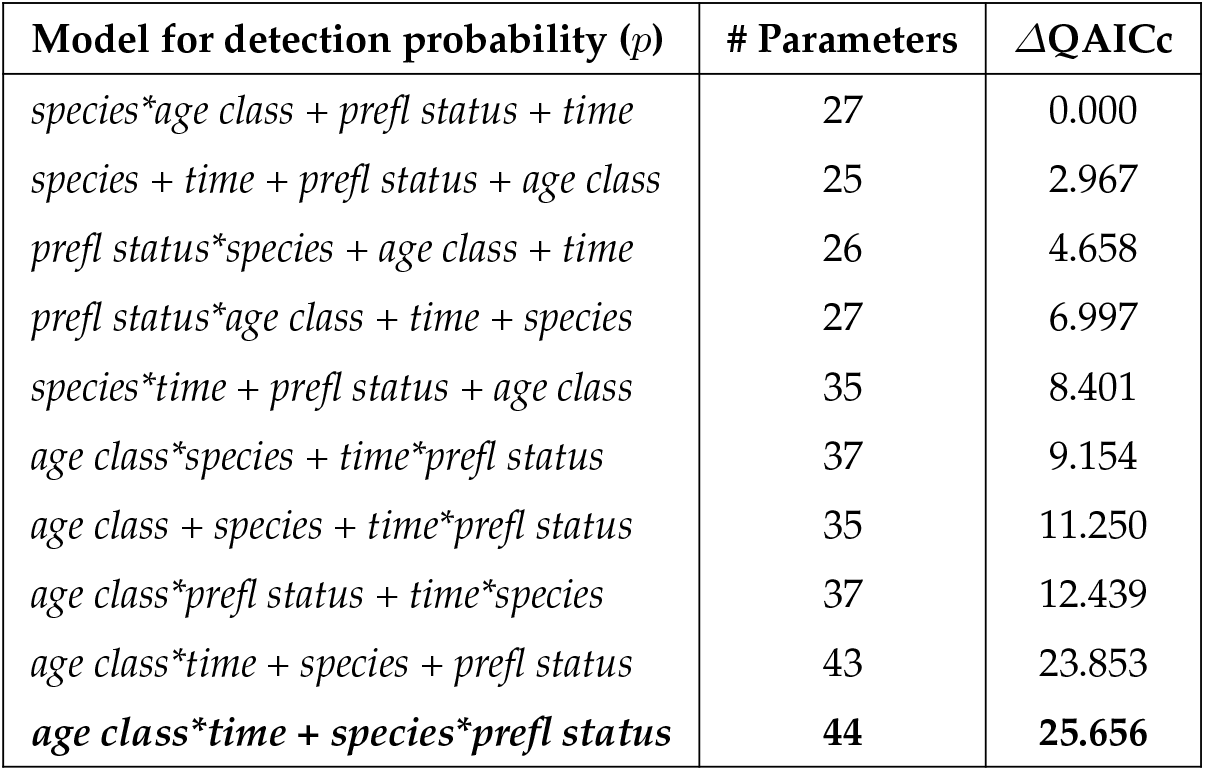
Candidate models for detection probability (*p*), each evaluated under the most complex model for survival probability (*Φ* = prefl status*species + prefl status*age class + species*age class). The most parameterized model (in bold) was used to estimate the overdispersion parameter (*ĉ*= 1.257) for QAICc adjustment using a parametric bootstrap GOF procedure.

**Table S4.**
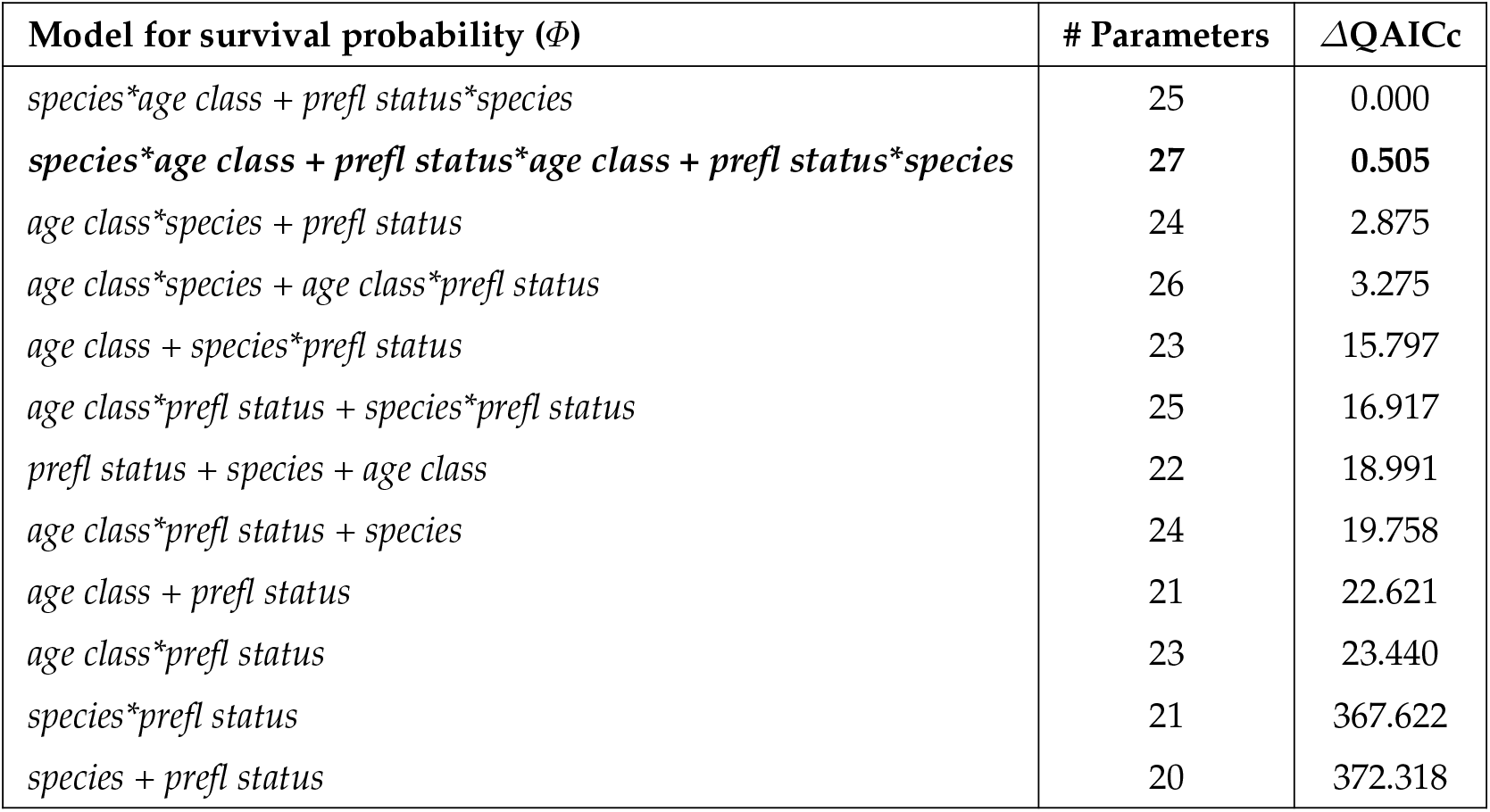
Candidate models for Survival probability (*Φ*), each evaluated with the best fitting model for detection probability (*p* = species*age class + prefl status + time). The most parameterized model (in bold) was used to estimate the overdispersion parameter (*ĉ*= 1.230) for QAICc adjustment using a parametric bootstrap GOF procedure.

**Table S5.**
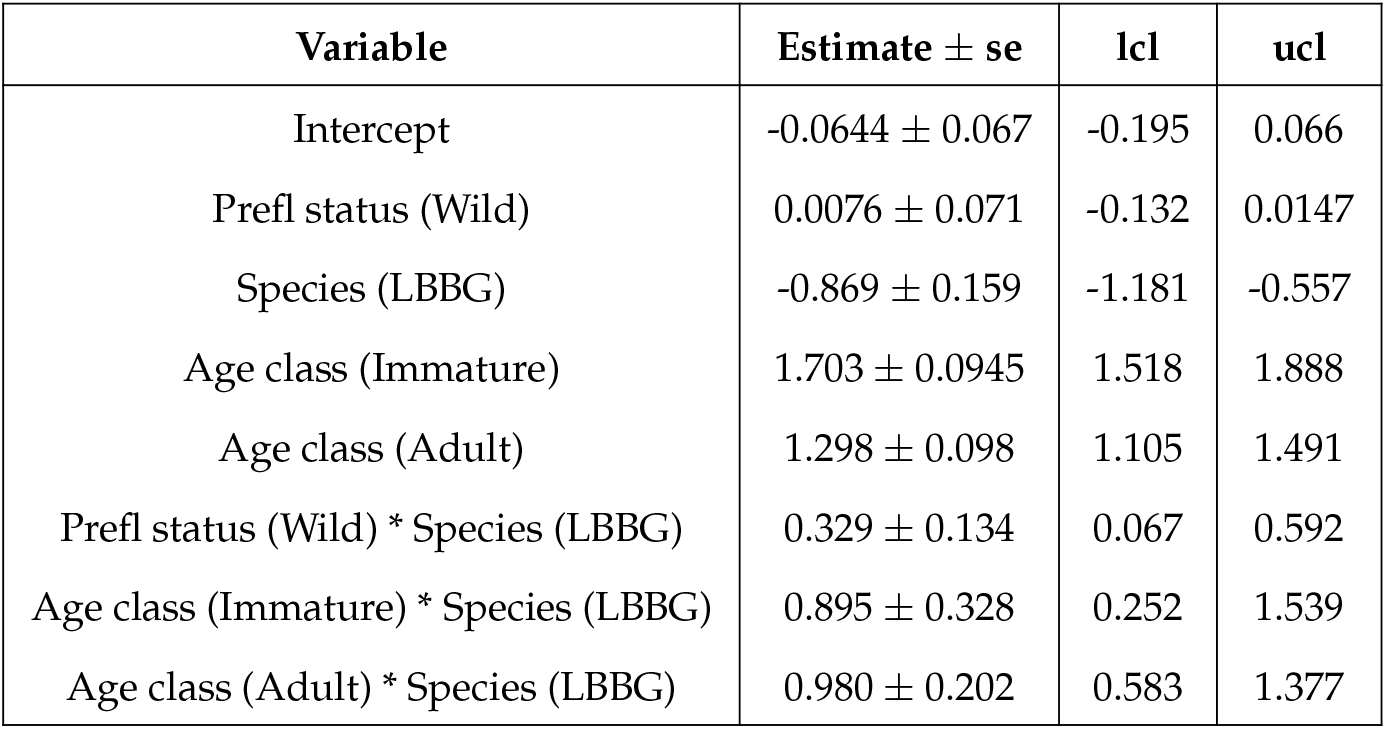
The output for the survival model species*age class + prefl status*species. Rehabilitated juvenile herring gulls are used as baseline (intercept).

| Variable | Estimate ± se | lcl | ucl |
| --- | --- | --- | --- |
| Intercept | -0.0644 ± 0.067 | -0.195 | 0.066 |
| Prefl status (Wild) | 0.0076 ± 0.071 | -0.132 | 0.0147 |
| Species (LBBG) | -0.869 ± 0.159 | -1.181 | -0.557 |
| Age class (Immature) | 1.703 ± 0.0945 | 1.518 | 1.888 |
| Age class (Adult) | 1.298 ± 0.098 | 1.105 | 1.491 |
| Prefl status (Wild) * Species (LBBG) | 0.329 ± 0.134 | 0.067 | 0.592 |
| Age class (Immature) * Species (LBBG) | 0.895 ± 0.328 | 0.252 | 1.539 |
| Age class (Adult) * Species (LBBG) | 0.980 ± 0.202 | 0.583 | 1.377 |

**Table S6.**
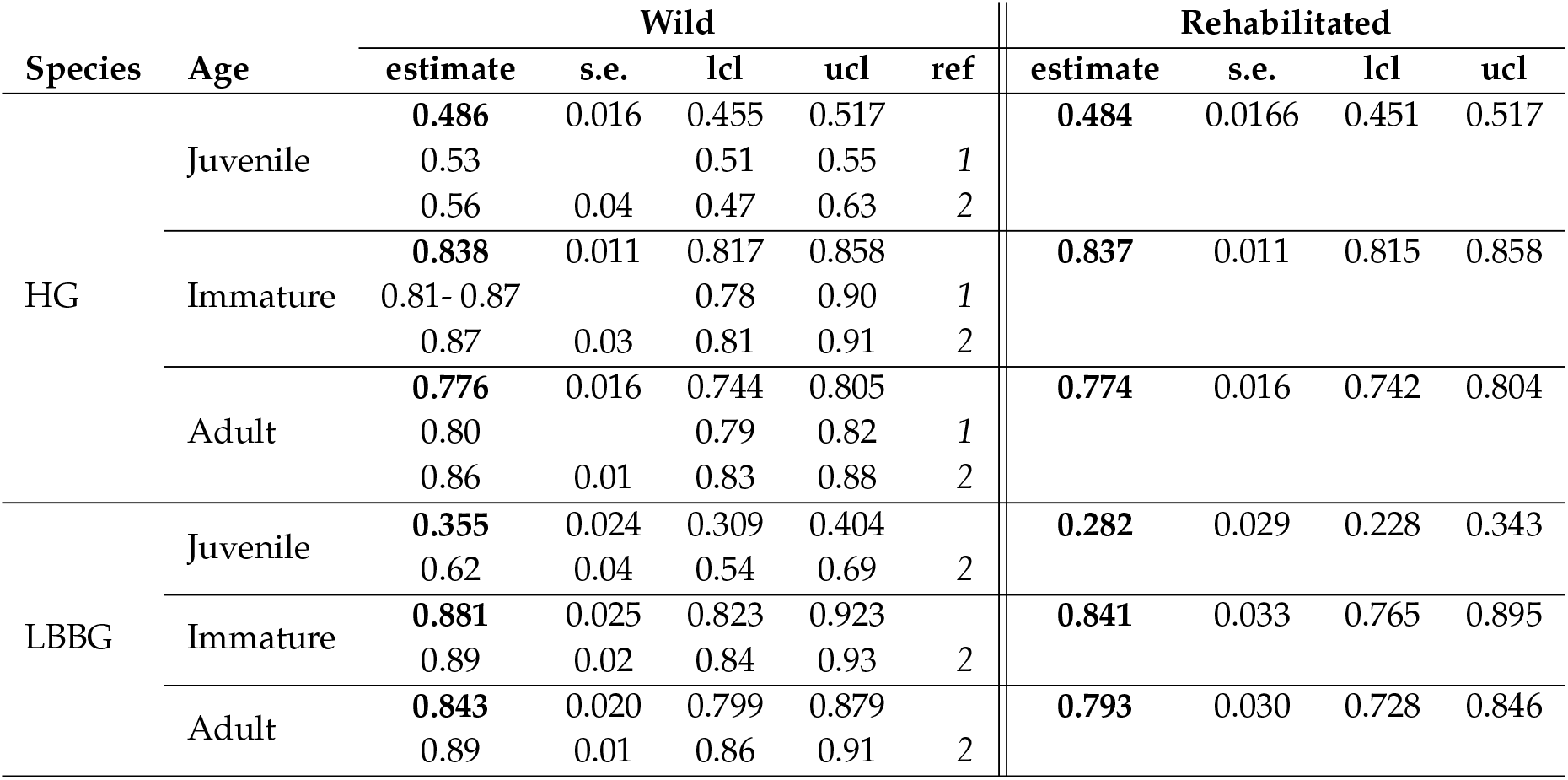
Survival estimates (*Φ*) for rehabilitated and wild herring and lesser black-backed gulls, in comparison with two other studies (*1*: Kentie et al., 2022, *2*: Schekkerman et al., 2021) who calculated survival rates for the same three age classes. Note that Schekkerman et al. (2021) included data from birds ringed as adults, which may limit the comparability of this age class with our results.

**Table S7.**
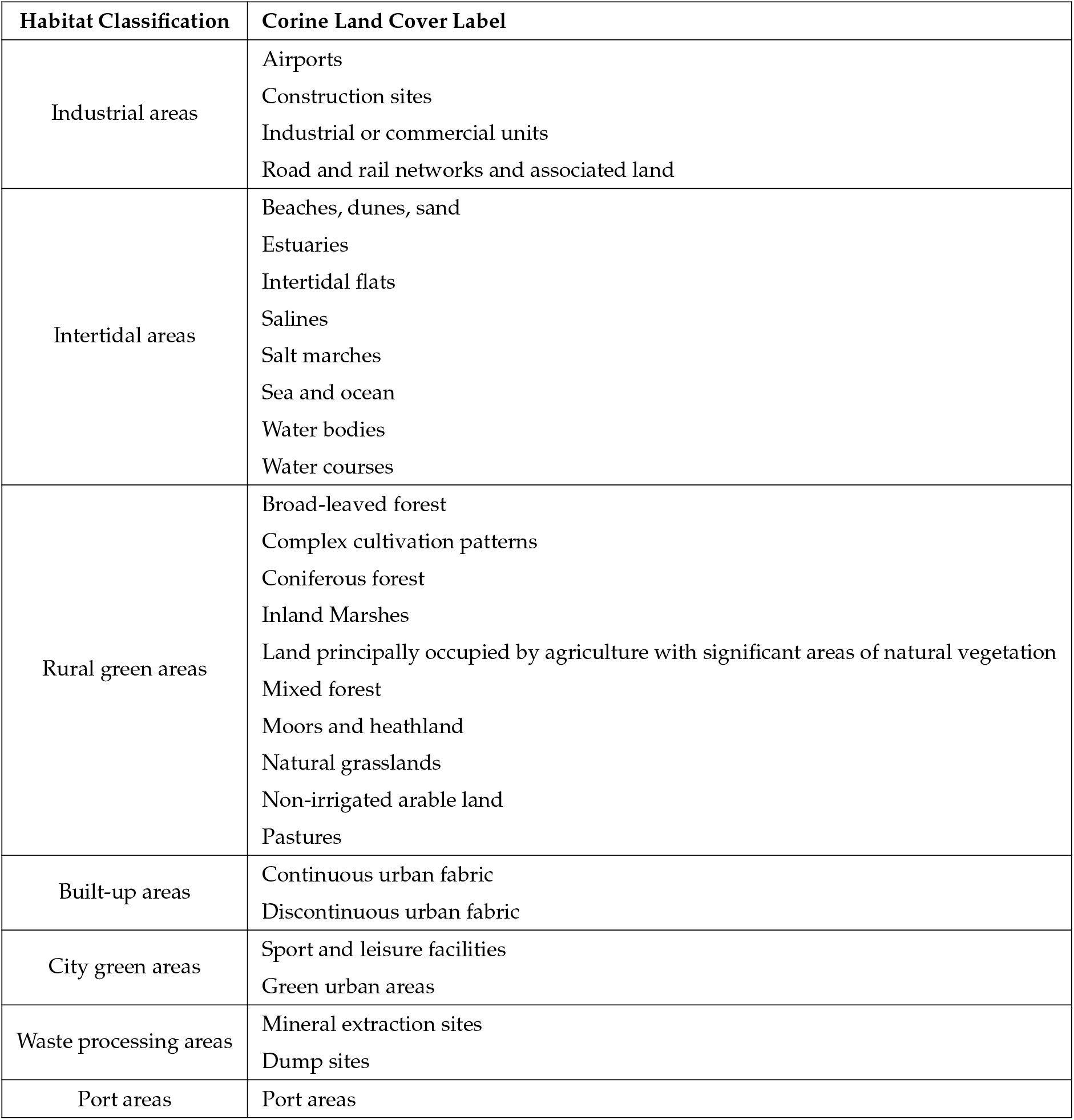
Habitat classification based on the 44 thematic classes of the Corine Land Cover inventory (100m, 2018).

**Table S8.**
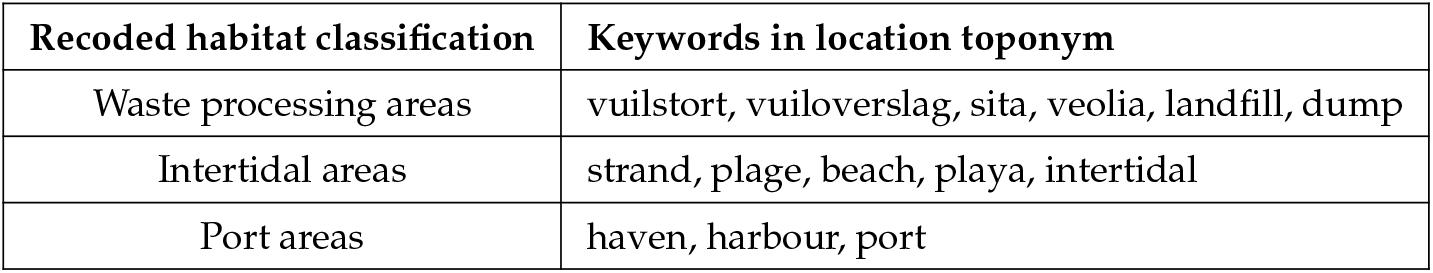
Manual recoding of habitat classifications based on location names to correct for imprecision in older GPS records. All other classifications were retained as originally annotated from the Corine Land Cover dataset.

**Table S9.**
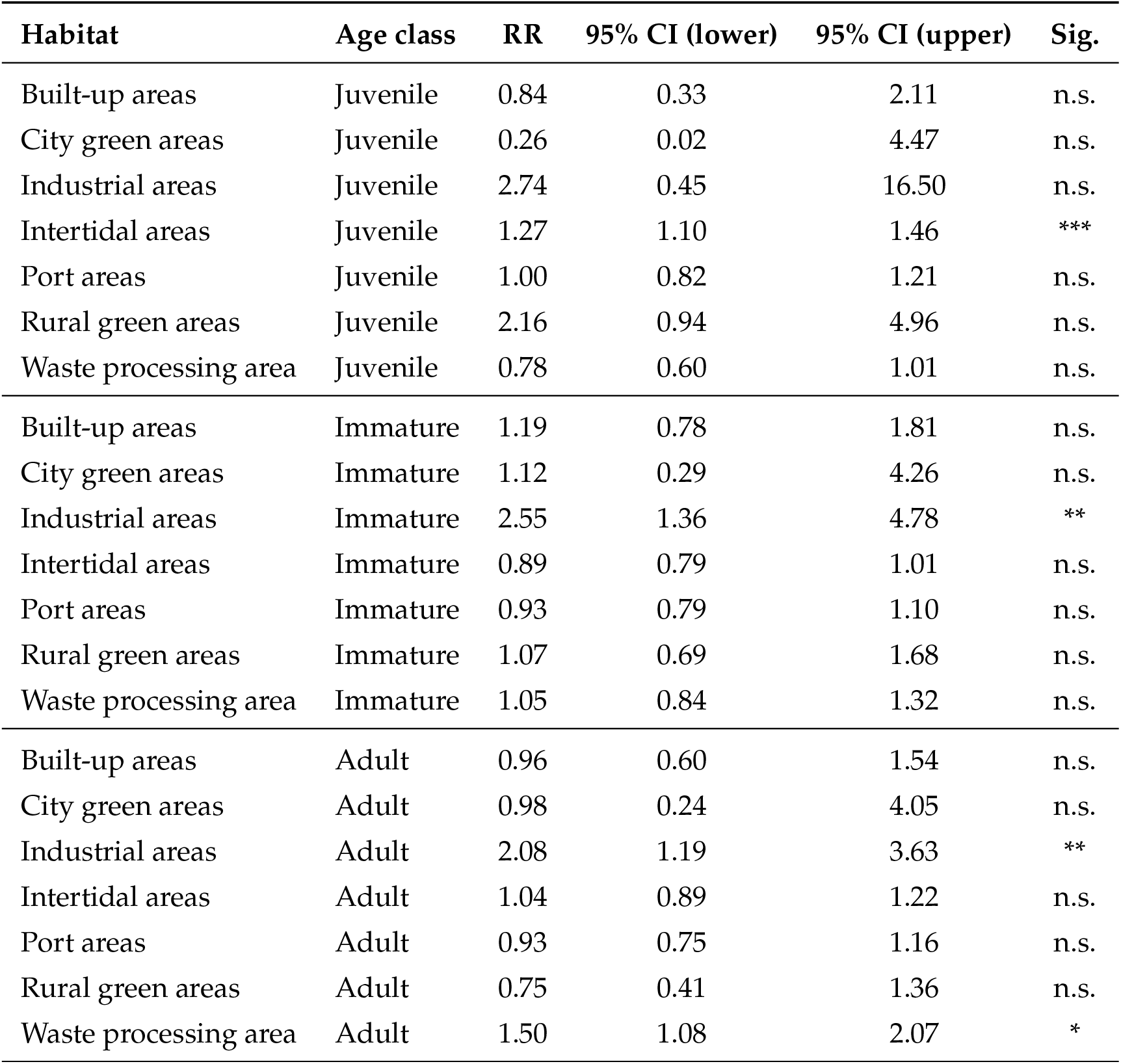
Habitat use contrasts (rehabilitated vs. wild) for herring gulls. Values show risk ratios (RR) with 95% confidence intervals. Significance: * *p <* 0.05, ** *p <* 0.01, *** *p <* 0.001, n.s. = not significant.

**Table S10.**
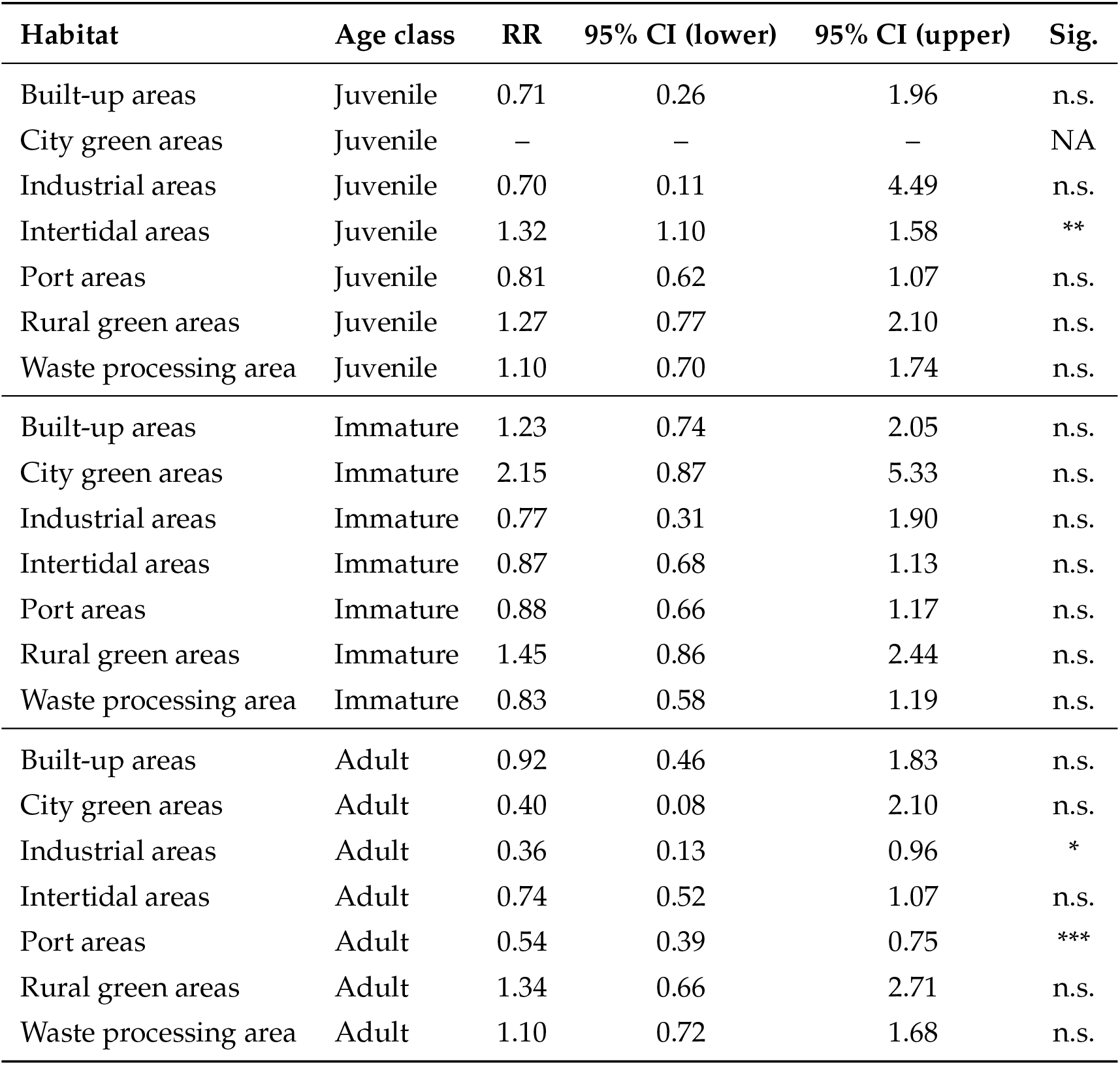
Habitat use contrasts (rehabilitated vs. wild) for lesser black-backed gulls. Values show risk ratios (RR) with 95% confidence intervals. Significance: * *p <* 0.05, ** *p <* 0.01, *** *p <* 0.001, n.s. = not significant, NA = not estimable.

**Table S11.**
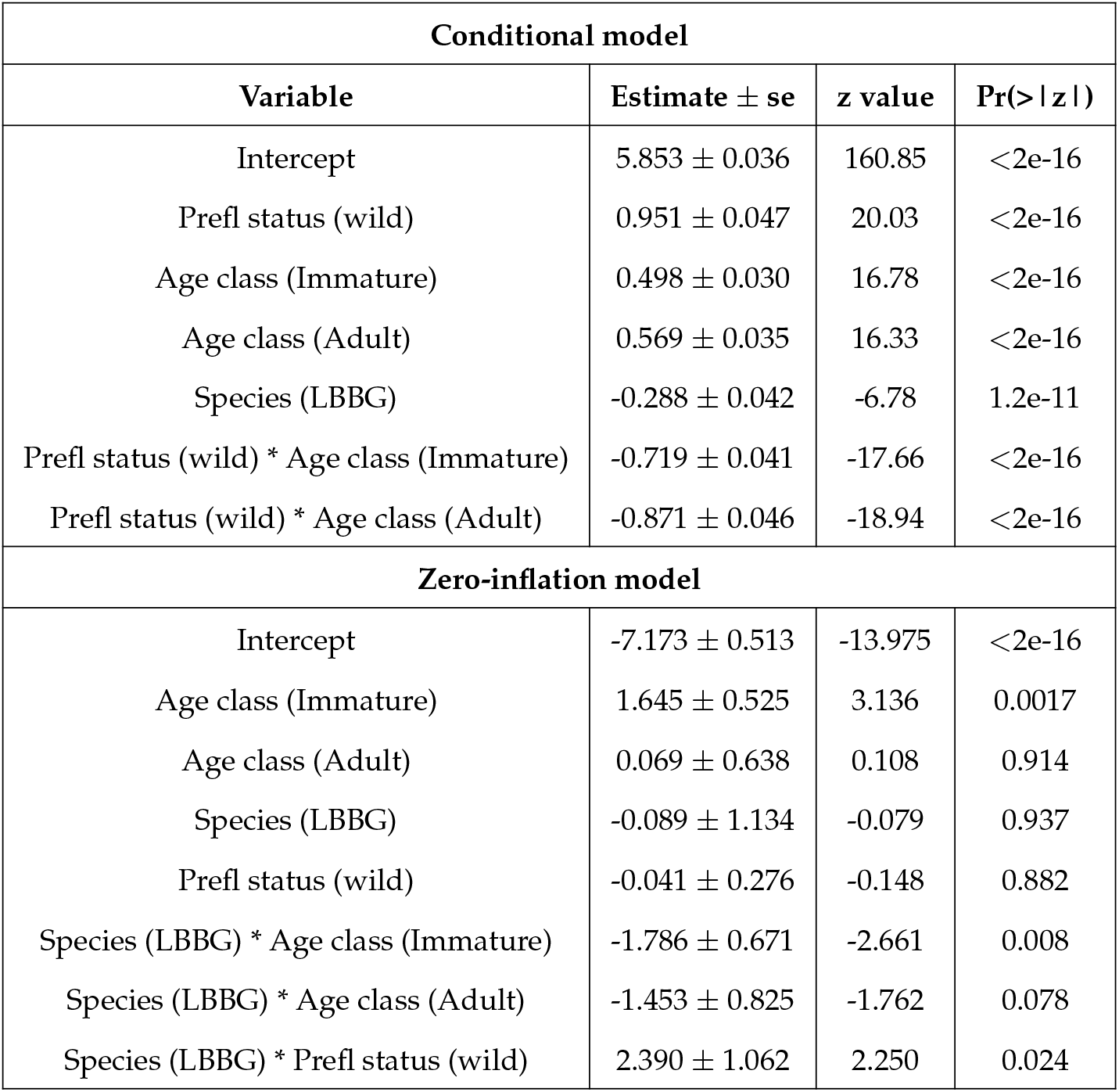
Model estimates for the conditional and zero-inflation parts of the zero-inflated Gaussian GLMM with log-transformed population density. Rehabilitated juvenile herring gulls are used as baseline (intercept).

| Conditional model |  |  |  |
| --- | --- | --- | --- |
| Variable | Estimate ± se | z value | Pr(> \|z\|) |
| Intercept | 5.853 ± 0.036 | 160.85 | <2e-16 |
| Prefl status (wild) | 0.951 ± 0.047 | 20.03 | <2e-16 |
| Age class (Immature) | 0.498 ± 0.030 | 16.78 | <2e-16 |
| Age class (Adult) | 0.569 ± 0.035 | 16.33 | <2e-16 |
| Species (LBBG) | -0.288 ± 0.042 | -6.78 | 1.2e-11 |
| Prefl status (wild) * Age class (Immature) | -0.719 ± 0.041 | -17.66 | <2e-16 |
| Prefl status (wild) * Age class (Adult) | -0.871 ± 0.046 | -18.94 | <2e-16 |
| Zero-inflation model |  |  |  |
| Intercept | -7.173 ± 0.513 | -13.975 | <2e-16 |
| Age class (Immature) | 1.645 ± 0.525 | 3.136 | 0.0017 |
| Age class (Adult) | 0.069 ± 0.638 | 0.108 | 0.914 |
| Species (LBBG) | -0.089 ± 1.134 | -0.079 | 0.937 |
| Prefl status (wild) | -0.041 ± 0.276 | -0.148 | 0.882 |
| Species (LBBG) * Age class (Immature) | -1.786 ± 0.671 | -2.661 | 0.008 |
| Species (LBBG) * Age class (Adult) | -1.453 ± 0.825 | -1.762 | 0.078 |
| Species (LBBG) * Prefl status (wild) | 2.390 ± 1.062 | 2.250 | 0.024 |

**Table S12.**
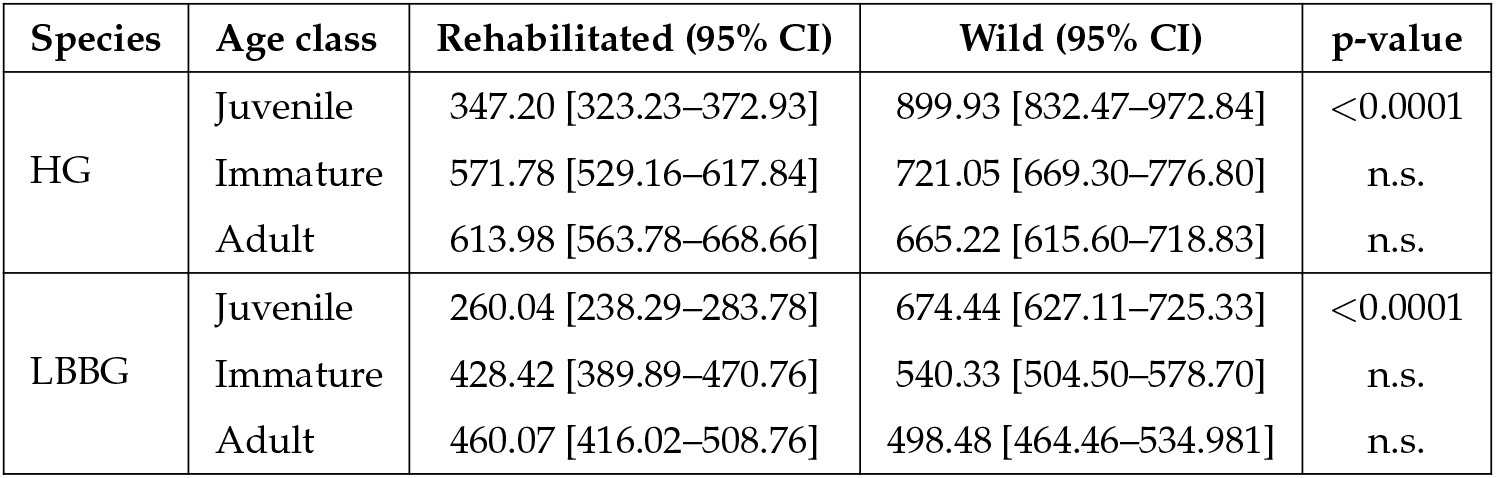
Predicted human population density (individuals per km^2^) at locations where rehabilitated and wild individuals were observed, derived from the conditional component of the zero-inflated GLMM. Values are model-based marginal means with 95% CIs (computed with `ggpredict()`, averaged over other covariates and random effects).P-values from Tukey-adjusted post hoc comparisons (Rehabilitated vs. Wild) within each species and age class are reported in the final column.

## Footnotes

1 R2ucare tests (Gimenez et al., 2018) detected significant trap-dependence across all groups; however because our analyses focus on relative rather than absolute survival, this is unlikely to affect the conclusions.

## References

Alden, B, Y Van Heezik, PJ Seddon, J Reid, and MJ Young (June 11, 2021). “Fat chance? Endangered penguin rehabilitation has mixed conservation outcomes”. In: Conservation Science and Practice 3.8, e452. DOI: 10.1111/csp2.452.

Allen, AM, BJ Ens, M van de Pol, R Visser, and PH Becker (2019). “Colour-ring wear and loss effects in citizen science mark-resighting studies”. In: Avian Research 10. Received 17 August 2018; accepted 08 April 2019; published 15 April 2019, p. 11. DOI: 10.1186/s40657-019-0151-z.

Anderson, DW, F Gress, and D Fry (1996). “Survival and dispersal of oiled brown pelicans after rehabilitation and release”. In: Marine Pollution Bulletin 32.10, pp. 711–718. ISSN: 0025-326X. DOI: 10.1016/0025-326X(96)00027-6.

Barker, RJ, KP Burnham, and GC White (2004). “Encounter history modeling of joint mark-recapture, tag-resighting and tag-recovery data under temporary emigration”. In: Statistica Sinica 14.4, pp. 1037–1055.

Blair, CD, LI Muller, JD Clark, and WH Stiver (Nov. 6, 2019). “Survival and Conflict Behavior of American Black Bears after Rehabilitation”. In: Journal of Wildlife Management 84.1, pp. 75–84. DOI: 10.1002/jwmg.21783.

Brooks, M, B Bolker, K Kristensen, M Maechler, A Magnusson, H Skaug, A Nielsen, C Berg, and K Van Bentham (Feb. 20, 2017). glmmTMB: Generalized Linear Mixed Models using Template Model Builder. DOI: 10.32614/cran.package.glmmtmb.

Burger, J (1981). “On Becoming Independent in Herring Gulls: Parent-Young Conflict”. In: The American Naturalist 117.4, pp. 444–456. ISSN: 00030147, 15375323.

Burnham, KP and DR Anderson (2002). Model Selection and Inference: A Practical Information-Theoretic Approach. 2nd. New York: Springer-Verlag. DOI: 10.1007/b97636.

Bustnes, JO, B Moe, M Helberg, and RA Phillips (2013). “Rapid long-distance migration in Norwegian Lesser Black-backed Gulls Larus fuscus fuscus along their eastern flyway”. In: Ibis 155.1, pp. 148–152. DOI: 10.1111/ibi.12022.

Camphuysen, CJ and A Gronert (2012). “Apparent Survival and Fecundity of Sympatric Lesser Black-Backed Gulls and Herring Gulls with Contrasting Population Trends”. In: Ardea 100.2, pp. 113–122. DOI: 10.5253/078.100.0202.

Chilvers, B, K Morgan, G Finlayson, and K Sievwright (2015). “Diving behaviour of wildlife impacted by an oil spill: A clean-up and rehabilitation success?” In: Marine Pollution Bulletin 100.1, pp. 128–133. ISSN: 0025-326X. DOI: 10.1016/j.marpolbul.2015.09.019.

Cope, HR, C McArthur, CR Dickman, TM Newsome, R Gray, and CA Herbert (Mar. 17, 2022). “A systematic review of factors affecting wildlife survival during rehabilitation and release”. In: PLoS ONE 17.3, e0265514. DOI: 10.1371/journal.pone.0265514.

Cormack, RM (Dec. 1, 1964). “Estimates of Survival from the Sighting of Marked Animals”. In: Biometrika 51.3/4, p. 429. DOI: 10.2307/2334149.

De La Cruz, S, J Takekawa, KA Spragens, J Yee, R Golightly, G Massey, L Henkel, S Larsen, and M Ziccardi (2013). “Post-release survival of surf scoters following an oil spill: an experimental approach to evaluating rehabilitation success”. In: Marine Pollution Bulletin 67.1-2. USGS Publications Warehouse, pp. 100–106. DOI: 10.1016/j.marpolbul.2012.11.027.

Duerr, R, D Jaques, B Selby, J Skoglund, and S Kosina (Sept. 2023). “Medical History and Postrelease Survival of Rehabilitated California Brown Pelicans Pelecanus occidentalis californicus, 2009–2019”. In: Marine Ornithology 51, pp. 157–168. DOI: 10.5038/2074-1235.51.2.1529.

Ellis, DH, GF Gee, SG Hereford, GH Olsen, TD Chisolm, JM Nicolich, KA Sullivan, NJ Thomas, M Nagendran, and JS Hatfield (Jan. 1, 2000). “Post-release survival of hand-reared and parent-reared Mississippi sandhill cranes”. In: Ornithological Applications 102.1, p. 104. DOI: 10.1650/0010-5422(2000)102.

Freed, LA, K Davis, LR Iwanowicz, and JT Anderson (2021). “Closely related gull species show contrasting foraging strategies in an urban environment”. In: Scientific Reports 11, p. 24485. DOI: 10.1038/s41598-021-02821-y.

Gaydos, J, L Vilchis, M Lance, S Jeffries, A Thomas, P Harner, V Greenwood, and M Ziccardi (Nov. 2012). “Postrelease movement of rehabilitated harbor seal (Phoca Vitulina Richardii) pups compared with cohort-matched wild seal pups”. In: Marine Mammal Science 29. DOI: 10.1111/mms.12002.

Gimenez, Olivier, Lebreton, Jean-Dominique, Choquet Rémi, Pradel, and Roger (2018). “R2ucare: An R package to perform goodness-of-fit tests for capture–recapture models”. In: Methods in Ecology and Evolution 9.7, pp. 1749–1754.

Golightly, RT, SH Newman, EN Craig, HR Carter, and JAK Mazet (2002). “Survival and Behavior of Western Gulls Following Exposure to Oil and Rehabilitation”. In: Wildlife Society Bulletin (1973-2006) 30.2, pp. 539–546. ISSN: 00917648, 19385463.

Goumas, M, CR Berkin, CW Rayner, and NJ Boogert (2024). “From the sea to the city: explaining gulls’ use of urban habitats”. In: Frontiers in Ecology and Evolution Volume 12 - 2024. ISSN: 2296-701X. DOI: 10.3389/fevo.2024.1256911.

Greig, SA, JC Coulson, and P Monaghan (1983). “Age-related differences in foraging success in the herring gull (Larus argentatus)”. In: Anim. Behav. 31, pp. 1237–1243. DOI: 10.1016/S0003-3472(83)80030-X.

Griffin, AS (Feb. 1, 2004). “Social learning about predators: a review and prospectus”. In: Learning Behavior 32.1, pp. 131–140. DOI: 10.3758/bf03196014.

Groom, CJ, K Warren, and PR Mawson (Mar. 6, 2017). “Survival and reintegration of rehabilitated Carnaby’s Cockatoos Zanda latirostris into wild flocks”. In: Bird Conservation International 28.1, pp. 86–99. DOI: 10.1017/s0959270916000642.

Grüebler, MU and B Naef-Daenzer (2010). “Survival benefits of post-fledging care: experimental approach to a critical part of avian reproductive strategies”. In: Journal of Animal Ecology 79.2, pp. 334–341. DOI: 10.1111/j.1365-2656.2009.01650.x. eprint: https://besjournals.onlinelibrary.wiley.com/doi/pdf/10.1111/j.1365-2656.2009.01650.x.

Hagen, CA, JM Goodell, BA Millsap, and GS Zimmerman (2024). “‘Dead birds flying’: can north American rehabilitated raptors released into the wild mitigate anthropogenic mortality?” In: Wildlife Biology 2024.5, e01283. DOI: 10.1002/wlb3.01283. eprint: https://nsojournals.onlinelibrary.wiley.com/doi/pdf/10.1002/wlb3.01283.

Hanson, M, N Hollingshead, K Schuler, WF Siemer, P Martin, and EM Bunting (Sept. 21, 2021). “Species, causes, and outcomes of wildlife rehabilitation in New York State”. In: PLoS ONE 16.9, e0257675. DOI: 10.1371/journal.pone.0257675.

Hartig, F (Aug. 26, 2016). DHARMa: Residual Diagnostics for Hierarchical (Multi-Level / Mixed) Regression Models. DOI: 10.32614/cran.package.dharma.

International Wildlife Rehabilitation Council (2024). URL: https://theiwrc.org/.

Inzani, E, L Kelley, R Thomas, and NJ Boogert (July 2024). “Early-life diet does not affect preference for fish in herring gulls (Larus argentatus)”. In: PeerJ 12. eCollection 2024, e17565. DOI: 10.7717/peerj.17565.

Jolly, GM (June 1, 1965). “Explicit Estimates from Capture-Recapture Data with Both Death and Immigration-Stochastic Model”. In: Biometrika 52.1/2, p. 225. DOI: 10.2307/2333826.

Kavelaars, MM, JM Baert, EWM Stienen, J Shamoun-Baranes, L Lens, and W Müller (2020). “Breeding habitat loss reveals limited foraging flexibility and increases foraging effort in a colonial breeding seabird”. In: Movement Ecology 8, p. 45. DOI: 10.1186/s40462-020-00231-9.

Kentie, R, J Shamoun-Baranes, AL Spaans, and K Camphuysen (July 28, 2022). “Spatial patterns in age- and colony-specific survival in a long-lived seabird across 14 contrasting colonies”. In: Ibis 165.1, pp. 82–95. DOI: 10.1111/ibi.13120.

Klaassen, RHG, BJ Ens, J Shamoun-Baranes, KM Exo, and F Bairlein (2012). “Migration strategy of a flight generalist, the Lesser Black-backed Gull Larus fuscus”. In: Behavioral Ecology 23.1, pp. 58–68. DOI: 10.1093/beheco/arr150.

Kreger, MD, JS Hatfield, I Estevez, GF Gee, and DA Clugston (Mar. 17, 2005). “The effects of captive rearing on the behavior of newly-released whooping cranes (Grus americana)”. In: Applied Animal Behaviour Science 93.1-2, pp. 165–178. DOI: 10.1016/j.applanim.2004.12.004.

Kwok, ABC, R Haering, SK Travers, and P Stathis (Sept. 10, 2021). “Trends in wildlife rehabilitation rescues and animal fate across a six-year period in New South Wales, Australia”. In: PLoS ONE 16.9, e0257209. DOI: 10.1371/journal.pone.0257209.

Laake, J (2013). RMark: An R Interface for Analysis of Capture-Recapture Data with MARK. AFSC Processed Rep. 2013-01. Seattle, WA: Alaska Fish. Sci. Cent., NOAA, Natl. Mar. Fish. Serv., p. 25.

Lamb, J, C Fiorello, Y Satgé, K Mills, M Ziccardi, and P Jodice (June 2018). “Movement patterns of California brown pelicans (Pelecanus occidentalis californicus) following oiling and rehabilitation”. In: Marine Pollution Bulletin 131, pp. 22–31. DOI: 10.1016/j.marpolbul.2018.03.043.

Lebreton, JD, K Burnham, J Clobert, and D Anderson (1992). “Modeling Survival and Testing Biological Hypotheses Using Marked Animals: A Unified Approach with Case Studies”. In: Ecological Monographs 62, pp. 67–118. DOI: 10.2307/2937171.

Lenth, RV (2025). emmeans: Estimated Marginal Means, aka Least-Squares Means. R package version 1.11.2-00002.

López-Idiáquez, D, P Vergara, JA Fargallo, and J Martínez-Padilla (2018). “Providing longer postfledging periods increases offspring survival at the expense of future fecundity”. In: PLoS One 13.9, e0203152. DOI: 10.1371/journal.pone.0203152.

Lüdecke, D, D Makowski, MS Ben-Shachar, I Patil, P Waggoner, BM Wiernik, and R Thériault (Apr. 24, 2019). performance: Assessment of Regression Models Performance. DOI: 10.32614/cran.package.performance.

Miller, TK, K Pierce, EE Clark, and RB Primack (2023). “Wildlife rehabilitation records reveal impacts of anthropogenic activities on wildlife health”. In: Biological Conservation 286, p. 110295. ISSN: 0006-3207. DOI: 10.1016/j.biocon.2023.110295.

Monadjem, A, K Wolter, W Neser, and A Kane (May 21, 2013). “Effect of rehabilitation on survival rates of endangered Cape vultures”. In: Animal Conservation 17.1, pp. 52–60. DOI: 10.1111/acv.12054.

Monaghan, P (1980). “Dominance and dispersal between feeding sites in the herring gull (Larus argentatus)”. In: Animal Behaviour 28.2, pp. 521–527. ISSN: 0003-3472. DOI: 10.1016/S0003-3472(80)80060-1.

Monaghan, P (1979). “Aspects of the breeding biology of Herring Gulls (Larus argentatus) in urban colonies”. In: Ibis 121.4, pp. 475–481. DOI: 10.1111/j.1474-919X.1979.tb06687.x.

Montesdeoca, N, P Calabuig, JA Corbera, JE Cooper, and J Orós (Oct. 2, 2017). “Causes of morbidity and mortality, and rehabilitation outcomes of birds in Gran Canaria Island, Spain”. In: Bird Study 64.4, pp. 523–534. DOI: 10.1080/00063657.2017.1411464.

Mullineaux, E and C Pawson (Dec. 26, 2023). “Trends in Admissions and Outcomes at a British Wildlife Rehabilitation Centre over a Ten-Year Period (2012–2022)”. In: Animals 14.1, p. 86. DOI: 10.3390/ani14010086.

Oppel, S, S Derville, P Pinet, UN Glutz von Blotzheim, and H Fritz (2015). “High juvenile mortality during migration in a declining population of a long-distance migratory raptor”. In: Ibis 157, pp. 545–557. DOI: 10.1111/ibi.12258.

Pesaresi, M, M Schiavina, P Politis, S Freire, K Krasnodębska, JH Uhl, A Carioli, C Corbane, L Dijkstra, P Florio, HK Friedrich, J Gao, S Leyk, L Lu, L Maffenini, I Mari-Rivero, M Melchiorri, V Syrris, J Van Den Hoek, and T Kemper (Aug. 30, 2024). “Advances on the global human settlement layer by joint assessment of earth observation and population survey data”. In: International Journal of Digital Earth 17.1. DOI: 10.1080/17538947.2024.2390454.

Punch, P (2001). “A retrospective study of the success of medical and surgical treatment of wild Australian raptors”. In: Australian Veterinary Journal 79.11, pp. 747–752. DOI: 10.1111/j.1751-0813.2001.tb10890.x. eprint: https://onlinelibrary.wiley.com/doi/pdf/10.1111/j.1751-0813.2001.tb10890.x.

Pyke, GH and JK Szabo (Mar. 23, 2017). “Conservation and the 4 Rs, which are rescue, rehabilitation, release, and research”. In: Conservation Biology 32.1, pp. 50–59. DOI: 10.1111/cobi.12937.

Reed, TE, LEB Kruuk, S Wanless, M Frederiksen, EJA Cunningham, and MP Harris (2008). “Reproductive Senescence in a Long-Lived Seabird: Rates of Decline in Late-Life Performance Are Associated with Varying Costs of Early Reproduction”. In: The American Naturalist 172.1, E20–E33. DOI: 10.1086/524957.

Rock, P (July 2005). “Urban gulls: Problems and solutions”. In: British Birds 98, pp. 338–355.

Rock, P and IP Vaughan (2013). “Long-term estimates of adult survival rates of urban Herring Gulls Larus argentatus and Lesser Black-backed Gulls Larus fuscus”. In: Ringing & Migration 28.1. Received 19 March 2013; accepted 26 April 2013; published online 05 July 2013, pp. 21–29. DOI: 10.1080/03078698.2013.811179.

Rotics, S, S Turjeman, M Kaatz, YS Resheff, D Zurell, N Sapir, U Eggers, A Flack, W Fiedler, F Jeltsch, M Wikelski, and R Nathan (2016). “The challenges of the first migration: movement and behaviour of juvenile vs. adult white storks with insights regarding juvenile mortality”. In: Journal of Animal Ecology 85, pp. 938–947. DOI: 10.1111/1365-2656.12525.

Sangster, S, M Haulena, C Nordstrom, and JK Gaydos (2021). “Interannual differences in postrelease movements of rehabilitated harbor seal pups (Phoca vitulina richardii) in the Salish Sea”. In: Marine Mammal Science 37.1, pp. 64–79. DOI: 10.1111/mms.12739. eprint: https://onlinelibrary.wiley.com/doi/pdf/10.1111/mms.12739.

Schekkerman, H, F Arts, RJ Buijs, W Courtens, T van Daele, R Fijn, A van Kleunen, H van der Jeugd, M Roodbergen, E Stienen, L de Vries, and B Ens (2021). Geïntegreerde populatieanalyse van vijf soorten kustbroedvogels in het Zuidwestelijk Deltagebied. Tech. rep. Sovon-rapport 2021/03, CAPSrapport 2021/01. Nijmegen: Sovon Vogelonderzoek Nederland.

Seber, GAF (June 1, 1965). “A Note on the Multiple-Recapture Census”. In: Biometrika 52.1/2, p. 249. DOI: 10.2307/2333827.

Service, CLM (2020). CORINE Land Cover 2018 (raster 100 m), Europe, 6-yearly - version 2020_2_0u1, May2020. DOI: 10.2909/960998c1-1870-4e82-8051-6485205ebbac.

Sharp, BE (1996). “Post-release survival of oiled, cleaned seabirds in North America”. In: Ibis 138.2, pp. 222–228. DOI: 10.1111/j.1474-919X.1996.tb04332.x. eprint: https://onlinelibrary.wiley.com/doi/pdf/10.1111/j.1474-919X.1996.tb04332.x.

Souëf, AL, C Holyoake, S Vitali, and K Warren (Feb. 3, 2015). “Presentation and prognostic indicators for free-living black cockatoos (Calyptorhynchus spp.) admitted to an Australian zoo veterinary hospital over 10 years”. In: Journal of Wildlife Diseases 51.2, pp. 380–388. DOI: 10.7589/2014-08-203.

Spelt, A, C Williamson, J Shamoun-Baranes, E Shepard, P Rock, and S Windsor (2019). “Habitat use of urban-nesting lesser black-backed gulls during the breeding season”. In: Scientific Reports 9.1, p. 10527. ISSN: 2045-2322. DOI: 10.1038/s41598-019-46890-6.

Stewart, LG, JL Lavers, ML Grant, PS Puskic, and AL Bond (2020). “Seasonal ingestion of anthropogenic debris in an urban population of gulls”. In: Marine Pollution Bulletin 160, p. 111549. DOI: 10.1016/j.marpolbul.2020.111549.

Sweeney, SJ, PT Redig, and HB Tordoff (1997). “Morbidity, survival and productivity of rehabilitated Peregrine Falcons in the upper midwestern US”. In: The journal of raptor research 31, pp. 347–352.

Thorup, K, F Korner-Nievergelt, EB Cohen, and SR Baillie (2014). “Large-scale spatial analysis of ringing and re-encounter data to infer movement patterns: A review including methodological perspectives”. In: Methods in Ecology and Evolution 5.12, pp. 1337–1350. DOI: 10.1111/2041-210X.12258. eprint: https://besjournals.onlinelibrary.wiley.com/doi/pdf/10.1111/2041-210X.12258.

Tribe, A and PR Brown (June 1, 2000). “The role of wildlife rescue groups in the care and rehabilitation of Australian fauna”. In: Human Dimensions of Wildlife 5.2, pp. 69–85. DOI: 10.1080/10871200009359180.

White, GC and KP Burnham (1999). “Program MARK: Survival estimation from populations of marked animals”. In: Bird Study 46. suppl, S120–S139. DOI: 10.1080/00063659909477239.

Zeileis, A, C Kleiber, and S Jackman (Jan. 1, 2008). “Regression Models for Count Data inR”. In: Journal of Statistical Software 27.8. DOI: 10.18637/jss.v027.i08.

