## Supplementary documents for "Evaluating wildlife rehabilitation: Successful return of rehabilitated gulls in the wild"

| Prefl cond | Species | 2012 | 2013 | 2014 | 2015 | 2016 | 2017 | 2018 | 2019 | Total |
|---|---|---|---|---|---|---|---|---|---|---|
| Rehabilitated | HG | 109 | 114 | 46 | 85 | 53 | 43 | 111 | 150 | **711** |
| | LBBG | 19 | 37 | 21 | 76 | 43 | 36 | 68 | 89 | **389** |
| Wild | HG | 119 | 21 | 79 | 28 | 139 | 294 | 77 | 69 | **826** |
| | LBBG | 316 | 20 | 184 | 89 | 199 | 327 | 170 | 212 | **1517** |

**Table S1.** The number of rehabilitated and wild juvenile herring and lesser black-backed gulls ringed each year (2012-2019)

| Model structure | | ΔQAICc | |
|---|---|---|---|
| **Survival probability ($\Phi$)** | **Detection probability ($p$)** | **HG ($\hat{c} = 1.290$)** | **LBBG ($\hat{c} = 1.181$)** |
| *prefl status + age class* | *age class + time + prefl status* | 0.000 | 0.000 |
| *prefl status*age class* | *age class + time + prefl status* | 1.748 | 3.313 |
| *prefl status + age class* | *age class*prefl status + time* | 3.918 | 3.458 |
| *prefl status*age class* | *age class*prefl status + time* | 5.556 | 5.617 |
| *prefl status + age class* | *age class + prefl status*time* | 12.224 | 8.587 |

**Table S2.** Top five ranked species-specific CJS models for herring gull (HG) and lesser black-backed gull (LBBG). Despite being modeled separately, the same structure ranked highest for both species, justifying the combined two-species analysis presented in the main text. $\hat{c}$ values: HG = 1.290, LBBG = 1.181.

| Model for detection probability ($p$) | # Parameters | ΔQAICc |
|---|---|---|
| *species*age class + prefl status + time* | 27 | 0.000 |
| *species + time + prefl status + age class* | 25 | 2.967 |
| *prefl status*species + age class + time* | 26 | 4.658 |
| *prefl status*age class + time + species* | 27 | 6.997 |
| *species*time + prefl status + age class* | 35 | 8.401 |
| *age class*species + time*prefl status* | 37 | 9.154 |
| *age class + species + time*prefl status* | 35 | 11.250 |
| *age class*prefl status + time*species* | 37 | 12.439 |
| *age class*time + species + prefl status* | 43 | 23.853 |
| ***age class*time + species*prefl status*** | **44** | **25.656** |

**Table S3.** Candidate models for detection probability ($p$), each evaluated under the most complex model for survival probability ($\Phi =$ prefl status*species + prefl status*age class + species*age class). The most parameterized model (in bold) was used to estimate the overdispersion parameter ($\hat{c} = 1.257$) for QAICc adjustment using a parametric bootstrap GOF procedure.

| **Model for survival probability ($\Phi$)** | **# Parameters** | **$\Delta$QAICc** |
|---|---|---|
| *species*age class + prefl status*species* | 25 | 0.000 |
| ***species*age class + prefl status*age class + prefl status*species*** | **27** | **0.505** |
| *age class*species + prefl status* | 24 | 2.875 |
| *age class*species + age class*prefl status* | 26 | 3.275 |
| *age class + species*prefl status* | 23 | 15.797 |
| *age class*prefl status + species*prefl status* | 25 | 16.917 |
| *prefl status + species + age class* | 22 | 18.991 |
| *age class*prefl status + species* | 24 | 19.758 |
| *age class + prefl status* | 21 | 22.621 |
| *age class*prefl status* | 23 | 23.440 |
| *species*prefl status* | 21 | 367.622 |
| *species + prefl status* | 20 | 372.318 |

**Table S4.** Candidate models for Survival probability ($\Phi$), each evaluated with the best fitting model for detection probability ($p =$ species*age class + prefl status + time). The most parameterized model (in bold) was used to estimate the overdispersion parameter ($\hat{c} = 1.230$) for QAICc adjustment using a parametric bootstrap GOF procedure.

| **Variable** | **Estimate $\pm$ se** | **lcl** | **ucl** |
|---|---|---|---|
| Intercept | -0.0644 $\pm$ 0.067 | -0.195 | 0.066 |
| Prefl status (Wild) | 0.0076 $\pm$ 0.071 | -0.132 | 0.0147 |
| Species (LBBG) | -0.869 $\pm$ 0.159 | -1.181 | -0.557 |
| Age class (Immature) | 1.703 $\pm$ 0.0945 | 1.518 | 1.888 |
| Age class (Adult) | 1.298 $\pm$ 0.098 | 1.105 | 1.491 |
| Prefl status (Wild) * Species (LBBG) | 0.329 $\pm$ 0.134 | 0.067 | 0.592 |
| Age class (Immature) * Species (LBBG) | 0.895 $\pm$ 0.328 | 0.252 | 1.539 |
| Age class (Adult) * Species (LBBG) | 0.980 $\pm$ 0.202 | 0.583 | 1.377 |

**Table S5.** The output for the survival model species*age class + prefl status*species. Rehabilitated juvenile herring gulls are used as baseline (intercept).

| Species | Age | Wild estimate | s.e. | lcl | ucl | ref | Rehabilitated estimate | s.e. | lcl | ucl |
|---|---|---|---|---|---|---|---|---|---|---|
| HG | Juvenile | **0.486** | 0.016 | 0.455 | 0.517 | | **0.484** | 0.0166 | 0.451 | 0.517 |
| | | 0.53 | | 0.51 | 0.55 | *1* | | | | |
| | | 0.56 | 0.04 | 0.47 | 0.63 | *2* | | | | |
| | Immature | **0.838** | 0.011 | 0.817 | 0.858 | | **0.837** | 0.011 | 0.815 | 0.858 |
| | | 0.81- 0.87 | | 0.78 | 0.90 | *1* | | | | |
| | | 0.87 | 0.03 | 0.81 | 0.91 | *2* | | | | |
| | Adult | **0.776** | 0.016 | 0.744 | 0.805 | | **0.774** | 0.016 | 0.742 | 0.804 |
| | | 0.80 | | 0.79 | 0.82 | *1* | | | | |
| | | 0.86 | 0.01 | 0.83 | 0.88 | *2* | | | | |
| LBBG | Juvenile | **0.355** | 0.024 | 0.309 | 0.404 | | **0.282** | 0.029 | 0.228 | 0.343 |
| | | 0.62 | 0.04 | 0.54 | 0.69 | *2* | | | | |
| | Immature | **0.881** | 0.025 | 0.823 | 0.923 | | **0.841** | 0.033 | 0.765 | 0.895 |
| | | 0.89 | 0.02 | 0.84 | 0.93 | *2* | | | | |
| | Adult | **0.843** | 0.020 | 0.799 | 0.879 | | **0.793** | 0.030 | 0.728 | 0.846 |
| | | 0.89 | 0.01 | 0.86 | 0.91 | *2* | | | | |

**Table S6.** Survival estimates ($\Phi$) for rehabilitated and wild herring and lesser black-backed gulls, in comparison with two other studies (*1*: Kentie et al., 2022, *2*: Schekkerman et al., 2021) who calculated survival rates for the same three age classes. Note that Schekkerman et al. (2021) included data from birds ringed as adults, which may limit the comparability of this age class with our results.

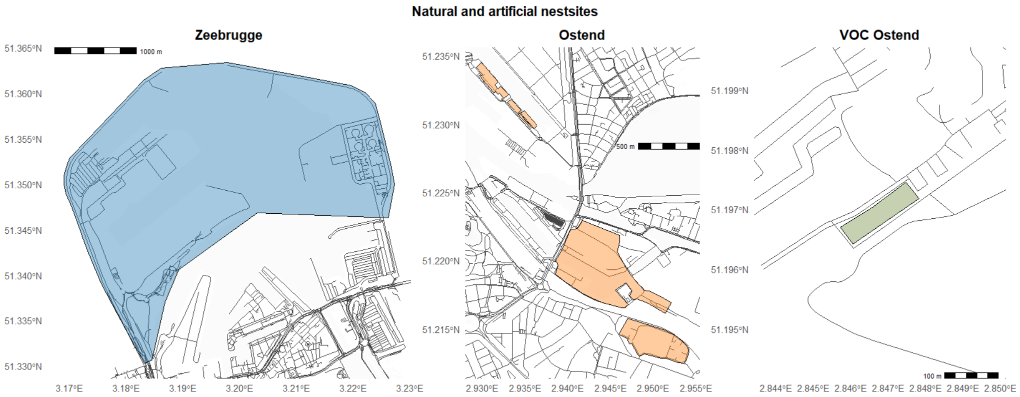


**Figure 5.** Natural nesting areas in Zeebrugge (left) and Ostend (right) and the rehabilitation centre as artificial nestsite.

| Habitat Classification | Corine Land Cover Label |
|---|---|
| Industrial areas | Airports<br>Construction sites<br>Industrial or commercial units<br>Road and rail networks and associated land |
| Intertidal areas | Beaches, dunes, sand<br>Estuaries<br>Intertidal flats<br>Salines<br>Salt marches<br>Sea and ocean<br>Water bodies<br>Water courses |
| Rural green areas | Broad-leaved forest<br>Complex cultivation patterns<br>Coniferous forest<br>Inland Marshes<br>Land principally occupied by agriculture with significant areas of natural vegetation<br>Mixed forest<br>Moors and heathland<br>Natural grasslands<br>Non-irrigated arable land<br>Pastures |
| Built-up areas | Continuous urban fabric<br>Discontinuous urban fabric |
| City green areas | Sport and leisure facilities<br>Green urban areas |
| Waste processing areas | Mineral extraction sites<br>Dump sites |
| Port areas | Port areas |

**Table S7.** Habitat classification based on the 44 thematic classes of the Corine Land Cover inventory (100m, 2018).

| Recoded habitat classification | Keywords in location toponym |
|---|---|
| Waste processing areas | vuilstort, vuiloverslag, sita, veolia, landfill, dump |
| Intertidal areas | strand, plage, beach, playa, intertidal |
| Port areas | haven, harbour, port |

**Table S8.** Manual recoding of habitat classifications based on location names to correct for imprecision in older GPS records. All other classifications were retained as originally annotated from the Corine Land Cover dataset.

| Habitat | Age class | RR | 95% CI (lower) | 95% CI (upper) | Sig. |
|---|---|---|---|---|---|
| Built-up areas | Juvenile | 0.84 | 0.33 | 2.11 | n.s. |
| City green areas | Juvenile | 0.26 | 0.02 | 4.47 | n.s. |
| Industrial areas | Juvenile | 2.74 | 0.45 | 16.50 | n.s. |
| Intertidal areas | Juvenile | 1.27 | 1.10 | 1.46 | *** |
| Port areas | Juvenile | 1.00 | 0.82 | 1.21 | n.s. |
| Rural green areas | Juvenile | 2.16 | 0.94 | 4.96 | n.s. |
| Waste processing area | Juvenile | 0.78 | 0.60 | 1.01 | n.s. |
| Built-up areas | Immature | 1.19 | 0.78 | 1.81 | n.s. |
| City green areas | Immature | 1.12 | 0.29 | 4.26 | n.s. |
| Industrial areas | Immature | 2.55 | 1.36 | 4.78 | ** |
| Intertidal areas | Immature | 0.89 | 0.79 | 1.01 | n.s. |
| Port areas | Immature | 0.93 | 0.79 | 1.10 | n.s. |
| Rural green areas | Immature | 1.07 | 0.69 | 1.68 | n.s. |
| Waste processing area | Immature | 1.05 | 0.84 | 1.32 | n.s. |
| Built-up areas | Adult | 0.96 | 0.60 | 1.54 | n.s. |
| City green areas | Adult | 0.98 | 0.24 | 4.05 | n.s. |
| Industrial areas | Adult | 2.08 | 1.19 | 3.63 | ** |
| Intertidal areas | Adult | 1.04 | 0.89 | 1.22 | n.s. |
| Port areas | Adult | 0.93 | 0.75 | 1.16 | n.s. |
| Rural green areas | Adult | 0.75 | 0.41 | 1.36 | n.s. |
| Waste processing area | Adult | 1.50 | 1.08 | 2.07 | * |

**Table S9.** Habitat use contrasts (rehabilitated vs. wild) for herring gulls. Values show risk ratios (RR) with 95% confidence intervals. Significance: * $p < 0.05$, ** $p < 0.01$, *** $p < 0.001$, n.s. = not significant.

| Habitat | Age class | RR | 95% CI (lower) | 95% CI (upper) | Sig. |
|---|---|---|---|---|---|
| Built-up areas | Juvenile | 0.71 | 0.26 | 1.96 | n.s. |
| City green areas | Juvenile | – | – | – | NA |
| Industrial areas | Juvenile | 0.70 | 0.11 | 4.49 | n.s. |
| Intertidal areas | Juvenile | 1.32 | 1.10 | 1.58 | ** |
| Port areas | Juvenile | 0.81 | 0.62 | 1.07 | n.s. |
| Rural green areas | Juvenile | 1.27 | 0.77 | 2.10 | n.s. |
| Waste processing area | Juvenile | 1.10 | 0.70 | 1.74 | n.s. |
| Built-up areas | Immature | 1.23 | 0.74 | 2.05 | n.s. |
| City green areas | Immature | 2.15 | 0.87 | 5.33 | n.s. |
| Industrial areas | Immature | 0.77 | 0.31 | 1.90 | n.s. |
| Intertidal areas | Immature | 0.87 | 0.68 | 1.13 | n.s. |
| Port areas | Immature | 0.88 | 0.66 | 1.17 | n.s. |
| Rural green areas | Immature | 1.45 | 0.86 | 2.44 | n.s. |
| Waste processing area | Immature | 0.83 | 0.58 | 1.19 | n.s. |
| Built-up areas | Adult | 0.92 | 0.46 | 1.83 | n.s. |
| City green areas | Adult | 0.40 | 0.08 | 2.10 | n.s. |
| Industrial areas | Adult | 0.36 | 0.13 | 0.96 | * |
| Intertidal areas | Adult | 0.74 | 0.52 | 1.07 | n.s. |
| Port areas | Adult | 0.54 | 0.39 | 0.75 | *** |
| Rural green areas | Adult | 1.34 | 0.66 | 2.71 | n.s. |
| Waste processing area | Adult | 1.10 | 0.72 | 1.68 | n.s. |

**Table S10.** Habitat use contrasts (rehabilitated vs. wild) for lesser black-backed gulls. Values show risk ratios (RR) with 95% confidence intervals. Significance: * $p < 0.05$, ** $p < 0.01$, *** $p < 0.001$, n.s. = not significant, NA = not estimable.

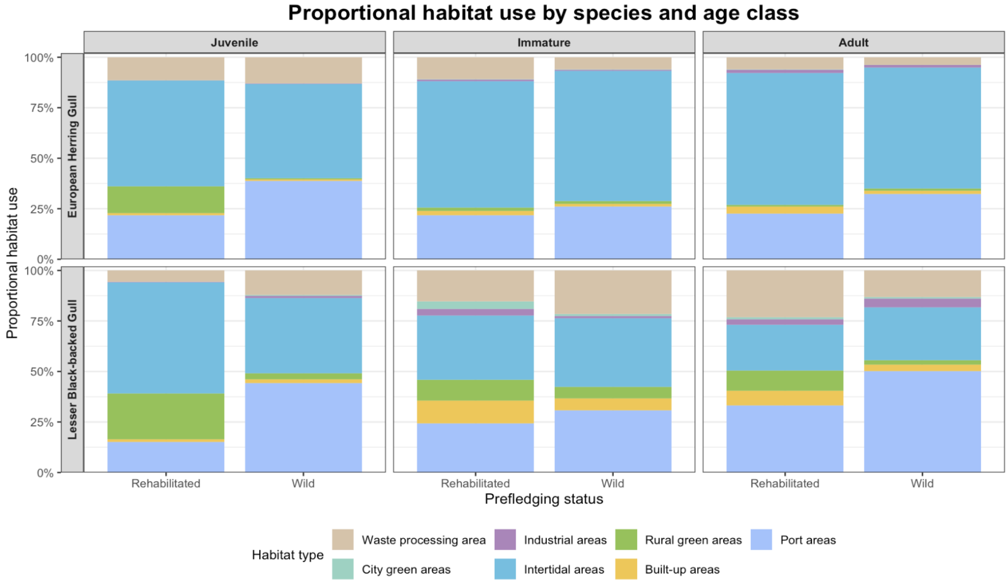


**Figure 6.** Proportional habitat use for herring gulls (top) and lesser black-backed gulls (bottom) across prefledging status and age classes. Proportions were calculated within each age class and prefledging status by normalizing predicted counts across habitat types

| Conditional model | | | |
|---|---|---|---|
| **Variable** | **Estimate ± se** | **z value** | **Pr(>\|z\|)** |
| Intercept | 5.853 ± 0.036 | 160.85 | <2e-16 |
| Prefl status (wild) | 0.951 ± 0.047 | 20.03 | <2e-16 |
| Age class (Immature) | 0.498 ± 0.030 | 16.78 | <2e-16 |
| Age class (Adult) | 0.569 ± 0.035 | 16.33 | <2e-16 |
| Species (LBBG) | -0.288 ± 0.042 | -6.78 | 1.2e-11 |
| Prefl status (wild) * Age class (Immature) | -0.719 ± 0.041 | -17.66 | <2e-16 |
| Prefl status (wild) * Age class (Adult) | -0.871 ± 0.046 | -18.94 | <2e-16 |
| **Zero-inflation model** | | | |
| Intercept | -7.173 ± 0.513 | -13.975 | <2e-16 |
| Age class (Immature) | 1.645 ± 0.525 | 3.136 | 0.0017 |
| Age class (Adult) | 0.069 ± 0.638 | 0.108 | 0.914 |
| Species (LBBG) | -0.089 ± 1.134 | -0.079 | 0.937 |
| Prefl status (wild) | -0.041 ± 0.276 | -0.148 | 0.882 |
| Species (LBBG) * Age class (Immature) | -1.786 ± 0.671 | -2.661 | 0.008 |
| Species (LBBG) * Age class (Adult) | -1.453 ± 0.825 | -1.762 | 0.078 |
| Species (LBBG) * Prefl status (wild) | 2.390 ± 1.062 | 2.250 | 0.024 |

**Table S11.** Model estimates for the conditional and zero-inflation parts of the zero-inflated Gaussian GLMM with log-transformed population density. Rehabilitated juvenile herring gulls are used as baseline (intercept).

| Species | Age class | Rehabilitated (95% CI) | Wild (95% CI) | p-value |
|---|---|---|---|---|
| HG | Juvenile | 347.20 [323.23–372.93] | 899.93 [832.47–972.84] | <0.0001 |
| | Immature | 571.78 [529.16–617.84] | 721.05 [669.30–776.80] | n.s. |
| | Adult | 613.98 [563.78–668.66] | 665.22 [615.60–718.83] | n.s. |
| LBBG | Juvenile | 260.04 [238.29–283.78] | 674.44 [627.11–725.33] | <0.0001 |
| | Immature | 428.42 [389.89–470.76] | 540.33 [504.50–578.70] | n.s. |
| | Adult | 460.07 [416.02–508.76] | 498.48 [464.46–534.981] | n.s. |

**Table S12.** Predicted human population density (individuals per $km^2$) at locations where rehabilitated and wild individuals were observed, derived from the conditional component of the zero-inflated GLMM. Values are model-based marginal means with 95% CIs (computed with `ggpredict()`, averaged over other covariates and random effects).P-values from Tukey-adjusted post hoc comparisons (Rehabilitated vs. Wild) within each species and age class are reported in the final column.
